# Asymmetric Depth Acclimation and Plasticity Limit the Refugial Potential of Mesophotic *Porites astreoides*

**DOI:** 10.64898/2026.03.18.712007

**Authors:** Elizaveta Skalon, Gretchen Goodbody-Gringley, Hagai Nativ, Shai Einbinder, Isabela Vitienes, Paul Zaslansky, Alex Chequer, Tali Mass

## Abstract

Mesophotic coral ecosystems have been proposed as climate refugia for shallow reefs, yet the capacity of mesophotic corals to persist across depth gradients remains unresolved. We conducted a 7-month reciprocal transplantation of the Caribbean coral *Porites astreoides* between shallow (10 m) and mesophotic (40 m) reefs to assess physiological, skeletal, and transcriptomic plasticity. Shallow colonies exhibited higher metabolic activity and skeletal extension, whereas mesophotic colonies showed reduced protein content, slower extension, and elevated expression of skeletal organic matrix genes. Transplant responses were asymmetric: shallow-to-deep corals acclimated through coordinated physiological and transcriptional adjustments, while deep-to-shallow transplants experienced elevated mortality and limited plasticity. SNP-based analyses suggest incomplete isolation across depth, while reciprocal transplant responses reflect greater phenotypic plasticity in shallow corals and indications of depth-specific specialization in mesophotic corals. Our findings indicate that shallow populations harbor greater acclimatory capacity, whereas mesophotic corals show constrained upward resilience, challenging the generality of deep reefs as refugia under rapid environmental change.

**Teaser:** Asymmetric plasticity limits the capacity of mesophotic corals to rejuvenate shallow reefs under climate change.

**Significance statement:** Mesophotic reefs have been proposed to support shallow coral persistence and recovery under ongoing climate change. Our study shows that this assumption can fail when populations have limited acclimatory capacity. In the reef-building coral *Porites astreoides*, individuals from shallow reefs tolerated translocation to mesophotic reefs better than mesophotic corals tolerated translocation upward, showing asymmetric phenotypic plasticity. Populations that persist within refugia can therefore have limited ability to support recovery elsewhere. At the same time, populations from more environmentally variable habitats may provide a source of tolerant individuals. The refugial potential of mesophotic reefs may thus depend on the interplay between phenotypic plasticity and environmental specialization across the depth gradient.

## Introduction

Coral reefs are among the most productive marine ecosystems in the world (1), largely due to the highly complex 3-dimensional calcium carbonate framework formed by corals and other calcifiers (2). However, coral cover has declined dramatically on reefs worldwide over the last 30 years (declining from ∼60% to <20% on some reefs), largely due to ocean warming (3–6). As shallow reefs continue to degrade, attention has increasingly turned to mesophotic coral ecosystems (MCEs; ∼30–150 m depth), because they experience reduced exposure to thermal extremes and wave-driven disturbance (7–9). These deeper habitats have therefore been proposed as potential climate refugia capable of buffering biodiversity loss and contributing to shallow reef recovery, a concept known as the Deep Reef Refugia Hypothesis (DRRH) (10, 11). However, the extent to which mesophotic corals can contribute to shallow-reef persistence depends on their physiological performance across depth gradients, their capacity for acclimation following environmental shifts, and the degree of connectivity between shallow and mesophotic populations, mediated by larval dispersal and allowing one habitat to serve as a source of recruits for the other (12).

Environmental conditions change predictably with depth. Light intensity (13), spectral composition (14), wave energy and temperature decline with depth (15). Corals spanning these gradients exhibit depth-associated differences in skeletal morphology, calcification, and photosynthetic performance (16–21). Such variation may arise through phenotypic plasticity, the ability of a genotype to produce different phenotypes across environments (22, 23), and through local genetic adaptation. Distinguishing between these mechanisms is essential for predicting vertical connectivity and resilience under climate change. Depth-generalist species, such as *Stylophora pistillata* in the Red Sea, and *Montastraea cavernosa* and *Porites astreoides* in the Western Atlantic, span broad bathymetric ranges (*13*, *19*, *21*, *24*, *25*), implying tolerance to environmental gradients and potential for gene flow across depths. In contrast, depth specialists such as *Alveopora spp*. in the Red Sea and *Leptoseris spp.* in the Caribbean (26, 27) may exhibit narrower ecological niches and stronger environmental specialization. Yet broad distributions alone do not resolve whether persistence across depths reflects individual plasticity or fixed population divergence. Integrating physiological performance with molecular responses is therefore necessary to evaluate the mechanisms underlying cross-depth persistence.

Theory and empirical evidence indicate that organisms originating from environmentally variable habitats evolve broader reaction norms and enhanced phenotypic plasticity (*28–30*). Repeated exposure to fluctuating stressors can favor mechanisms such as pre-activation of stress-response pathways, enhanced DNA repair capacity, and modulation of oxidative stress and metabolic processes, enabling rapid acclimation to novel environmental conditions (*29*, *31*, *32*). Shallow reefs experience greater fluctuations in irradiance, temperature, and hydrodynamics relative to mesophotic environments, potentially selecting for greater metabolic flexibility and enhanced stress tolerance. Conversely, the comparatively stable low-light conditions of mesophotic reefs may favor phenotypes optimized for depth-specific performance but less capable of rapid upward acclimation. If shallow and mesophotic populations differ in their capacity for phenotypic plasticity, resulting in asymmetric acclimation across depth gradients, this would have direct implications for the refugial capacity of deep reefs.

Here, we conducted a 7-month reciprocal transplantation of the Caribbean coral *Porites astreoides* between shallow (10 m) and mesophotic (40 m) reefs in Little Cayman. This period covered both winter and summer seasons in the Caribbean (*33*). By integrating skeletal extension measurements, physiological profiling, transcriptomic analyses, and SNP-based assessment of sample relatedness, we addressed three questions: (i) How strongly do season and depth affect coral performance? (ii) To what extent are depth-associated differences genetically determined or plastically regulated? (iii) Does the environment of origin influence coral acclimation across depth gradients?

By coupling ecological performance with transcriptomic data, we evaluate whether vertical persistence in *P. astreoides* is mediated primarily by phenotypic plasticity and assess the extent to which mesophotic populations can contribute to shallow reef resilience under ongoing environmental change.

## Results

### Study sites and environmental conditions

The study was conducted at two reef sites on Little Cayman Island, Martha’s Finyard (MF) and Coral City (CC) (Fig. 1A), which exhibit similar geomorphology and temperature conditions. At each site, colonies of *P. astreoides* were collected from shallow (10 m) and mesophotic (40 m) depths in winter, representing the origin groups S and D, respectively. Colonies were divided and returned to the reef in a reciprocal transplantation design with four treatments: shallow→shallow (SS), shallow→deep (SD), deep→deep (DD), and deep→shallow (DS), and were retrieved the following summer after seven months of experiment (Fig. S1).

**Figure 1.**
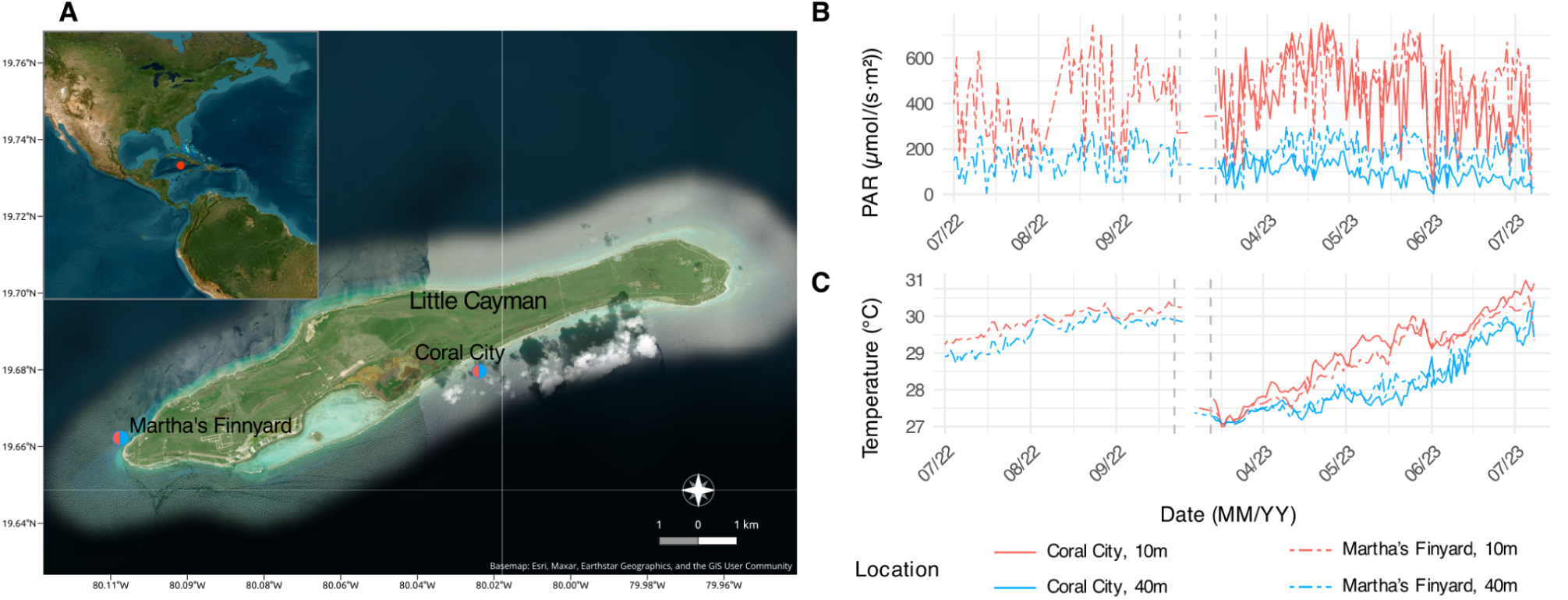
Environmental conditions and study sites in Little Cayman. **(A)** Map of study sites in Little Cayman. **(B)** Midday average of photosynthetically active radiation (PAR; μmol m^-2^ s^-1^), and **(C)** Daily average temperature (°C) from July 2022 to July 2023.

Midday photosynthetically active radiation (PAR; 12:00–14:00) measured between July 2022 and July 2023 averaged 447 ± 12 and 435 ± 16 μmol m^-^² s^-^¹ at 10 m and 171 ± 5 and 93 ± 4 μmol m^-^² s^-^¹ at 40 m at Martha’s Finyard and Coral City, respectively (mean ± SE; Fig. 1B, Table S1). PAR variability was several-fold greater in shallow than in mesophotic habitats at both sites. By contrast, seawater temperatures showed similar ranges across depths, varying between 27.0–31.0 °C at 10 m and 27.05–30.42 °C at 40 m. Monthly temperature fluctuations were comparable between depths and sites (∼1 °C), with maximum temperatures increasing from March to July 2023 at both depths (Fig. 1C, Table S2).

### Transplant survival of *P. astreoides*

Survival of *P. astreoides* after 7-month reciprocal translocation varied across sites and treatments (Fig. S2). Transplantation of shallow-origin corals to mesophotic depths had little effect on survival: SD colonies showed survival comparable to SS corals at both sites (100% at Coral City and 70% at Martha’s Finyard). In contrast, transplantation of mesophotic-origin corals to shallow reefs strongly reduced survival. DS colonies exhibited substantially lower survival than DD, particularly at Martha’s Finyard where survival dropped to 14.3% compared with 85.7% in DD colonies. A site-adjusted logistic regression indicated significant effects of both treatment and site on mortality, with DS transplants exhibiting significantly higher mortality than SS, SD, and DD treatments.

### Genet identity and relatedness across depths and sites

We used transcriptome-derived SNPs to explore genetic relatedness among sampled colonies. After collapsing replicates generated by experimental fragmentation of the same source colony (n = 20, 6,337 SNPs), KING kinship analysis showed that four of five shallow Martha’s Finyard (S.MF) samples and two of five shallow Coral City (S.CC) samples were identified as clonemates (pairwise kinship > 0.354), and additional first-degree relationships (pairwise kinship > 0.177) were detected within S.MF and among S.MF, deep Martha’s Finyard (D.MF) samples, and deep Coral City (D.CC) samples (Supplementary Fig. S3).

After clone/relative filtering (n=13, 6,337 SNPs), Fst estimates indicated little differentiation between mesophotic colonies from the two sites (Fst D.MF–D.CC = 0.01) and greater differentiation between shallow and mesophotic samples (Fst D.MF–S.CC = 0.10; Fst D.CC–S.CC = 0.06) (Supplementary Fig. S3).

### Variation in symbiont composition across experimental groups

Symbiont species identification revealed that corals from all groups hosted a similar consortium of Symbiodiniaceae, dominated by *Symbiodinium microadriaticum* and *S. linucheae* (Fig. S4). Permutational multivariate analysis of variance (PERMANOVA) indicated no significant differences in symbiont composition between experimental groups at either species level (R^2^ = 0.23, F = 0.59, p = 0.85; 10,000 permutations) or genus level (R² = 0.09, F = 0.50, p = 0.79; 10,000 permutations). Within- and between-group variations were also comparable at the species level (Wilcoxon test, W = 56, p = 0.25).

### Skeletal structure across treatments

The thick alizarin band indicates that *P. astreoides* tissue extends approximately 1.166 ± 0.04 mm (mean ± SE) into the skeleton in both mesophotic and shallow colonies (Fig. 2A) as previously reported for *Porites* (*34*). Skeletal linear extension differed significantly between translocation treatments (Fig. 2B). SS corals exhibited the highest extension rates (0.436 ± 0.02 cm year^-1^), whereas both SD transplants and DD corals showed significantly lower extension (0.35 ± 0.01 and 0.328 ± 0.01 cm year^-1^ respectively), with no significant difference between these two groups. DS transplants displayed intermediate extension rates (0.366 ± 0.02 cm year^-1^), which were higher than those of DD colonies but remained significantly lower than SS corals. No significant differences in corallite diameter or polyp density were detected among treatments (Fig. S5).

**Figure 2.**
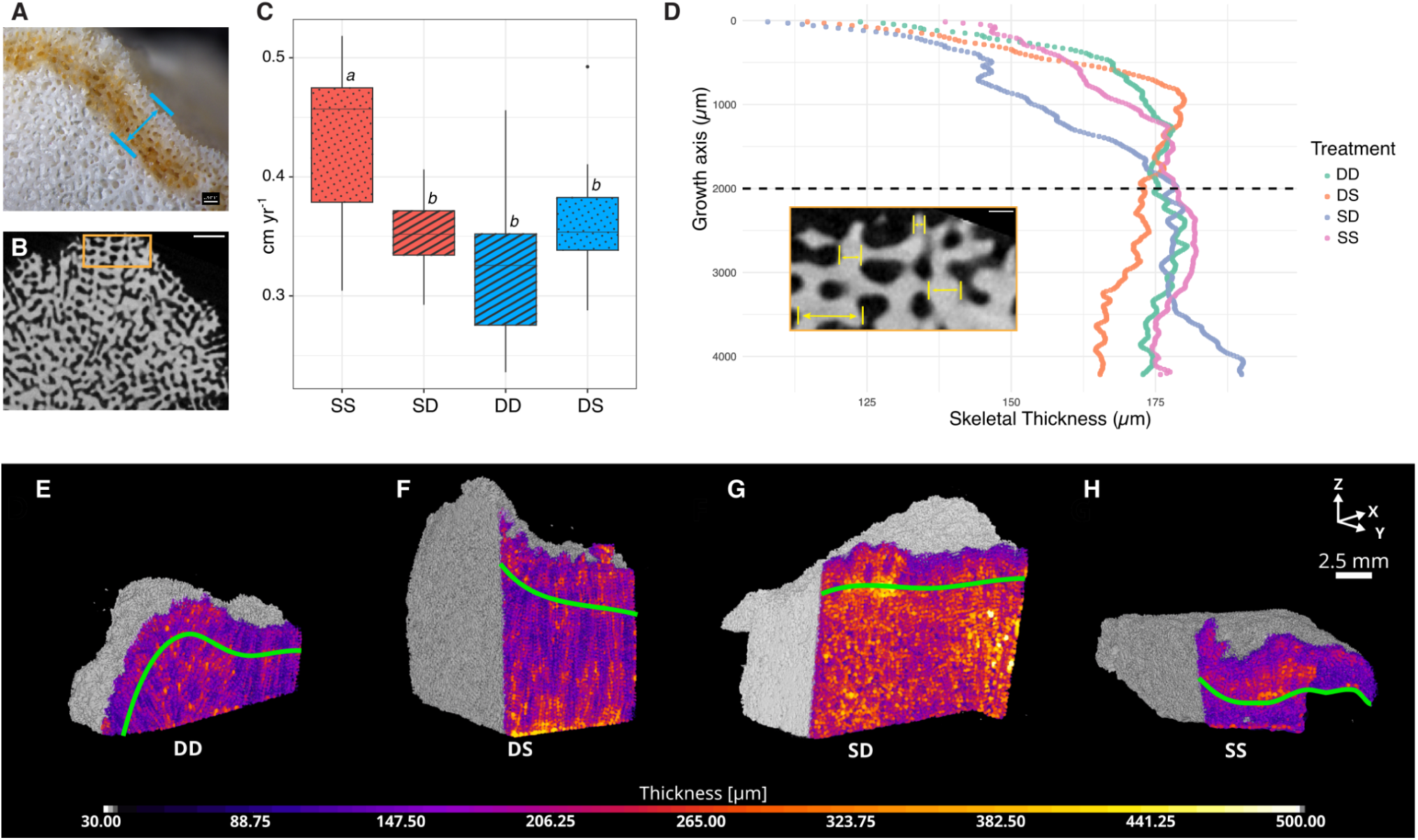
Skeletal growth and thickness following reciprocal transplantation. (A) Representative skeleton showing the alizarin-stained growth band marking skeletal accretion during the experimental period. (B) Longitudinal micro-CT section highlighting newly formed skeletal material. (C) Linear extension rates derived from the distance between the alizarin stain band and the skeletal surface (blue arrow in A). Different letters indicate significant differences between treatments (adjusted p < 0.05). (D) Mean skeletal thickness profiles along the growth axis (slice thickness = 15 μm). The dashed line marks the extent of new skeletal growth during transplantation (inset demonstrates the thickness measurement of the skeleton area). (E-H) Three-dimensional renderings of internal skeletal thickness for DD (E), DS (F), SD (G), and SS (H) treatments. All reconstructions use the same color scale for direct comparison. Green lines indicate the alizarin stain band.

Volumetric analysis across the growth axis further demonstrated that variation in linear extension was accompanied by changes in internal skeletal thickening (Fig. 2C). Mean skeletal thickness varied systematically along the growth axis, with pronounced differences across the skeletal extension region formed during the transplantation period, as indicated by the dashed line (Fig. 2D). Sliced top-view 3D renderings revealed the underlying skeletal mineral structure associated with these patterns (Fig. 2E-H), showing distinct internal thickness distributions in corals originating from different treatments.

### Coral physiological trait profiles by depth and season

#### Protein content and symbiont characteristics

Total protein content differed significantly between depths (Fig. S6, Table S3). Protein levels were considerably higher in shallow corals compared to mesophotic ones, both before translocation (S vs. D) and in corals maintained at their native depth after translocation (SS vs. DD). Cross-depth transplantation resulted in a dramatic decrease in total protein in SD corals (SD vs. DD), and, conversely, an increase in DS corals (DS vs. SS), indicating a strong dependence of protein content on depth. A significantly lower density of Symbiodiniaceae cells per cm^2^ was detected in DS corals compared with all other experimental groups except DD. Chlorophyll concentration per symbiont cell showed only mild season-dependent variation, with lower values observed before translocation and higher values in cross-depth translocated corals.

#### Photophysiological parameters

Across all photophysiological parameters, *P. astreoides* showed significant variation across depths only in photosynthetic efficiency (Fv/Fm), and the pattern was similar to changes in protein concentration. In winter, Fv/Fm did not differ between S and D corals. However, in summer, photosynthetic efficiency was significantly higher in DD colonies compared to SS colonies, and translocated corals shifted toward the values of the native population at their destination depth. Functional absorption cross-section (σPSII) and connectivity parameter (p) showed only seasonal dependence, with significantly higher values in summer compared to winter. The maximum photosynthesis rate (Pmax, s^-1^ PSII^-1^) was elevated in SS and SD corals compared to D corals, but showed no other significant differences.

#### Multivariate structure of physiological traits

A PCA of coral physiological traits pre- and post-translocation revealed separation between experimental groups (Fig. 3A). On both axes, Pmax was positively associated with σPSII and connectivity parameter (p). On PC1, they were also positively correlated to chlorophyll content per symbiont cell, contrasting with symbiont cell density, protein content, and photosynthetic efficiency, whereas on PC2 the direction of this correlation was reversed, and Fv/Fm showed little contribution to this axis (Fig. 3A, S7A). PAR was correlated with the ordination and mainly aligned with PC1 (R² = 0.23, p = 0.001), whereas temperature showed a weaker association, primarily oriented along PC2 (R² = 0.12, p = 0.005) (Table S4). Treatment had a significant effect on physiological trait composition (PERMANOVA, F= 7.95, R^2^ = 0.314, p = 0.001) (Table S5). Pairwise comparisons showed significant differences among most experimental groups (adjusted p ≤ 0.05; Table S6), with non-significant differences only among DD, SD, and DS, and between DS and SS, with separation being dominated by photophysiological traits. An additional PCA of coral physiological and skeletal traits included only corals post-transplantation (Fig. 3B, S7B). In this analysis, chlorophyll content per symbiont cell was negatively associated with symbiont cell density and protein content on both axes, consistent with the pattern observed in the previous PCA. On PC1, chlorophyll content per cell was also negatively associated with σPSII, the connectivity parameter (p), and photosynthetic efficiency, but positively correlated with Pmax. On PC2 the direction of these relationships was reversed. PAR (R^2^ = 0.57, p = 0.001) and temperature (R^2^ = 0.31, p = 0.002) were correlated with PC2 (Table S4). On the PCA plot SS treatment was clearly separated from DD treatment by higher values for skeletal linear extension and lower for photosynthetic efficiency, while cross-depth transplants occupied intermediate positions leaning toward DD colonies. PERMANOVA using an origin × transplant depth design detected significant effects of origin depth (p = 0.001), transplant depth (p = 0.001), and their interaction (p = 0.021) on trait composition, together explaining 20.5% of the multivariate variation (Table S5). Pairwise treatment comparisons further supported significant differences between SS and DD and SS and SD (Table S6).

**Figure 3.**
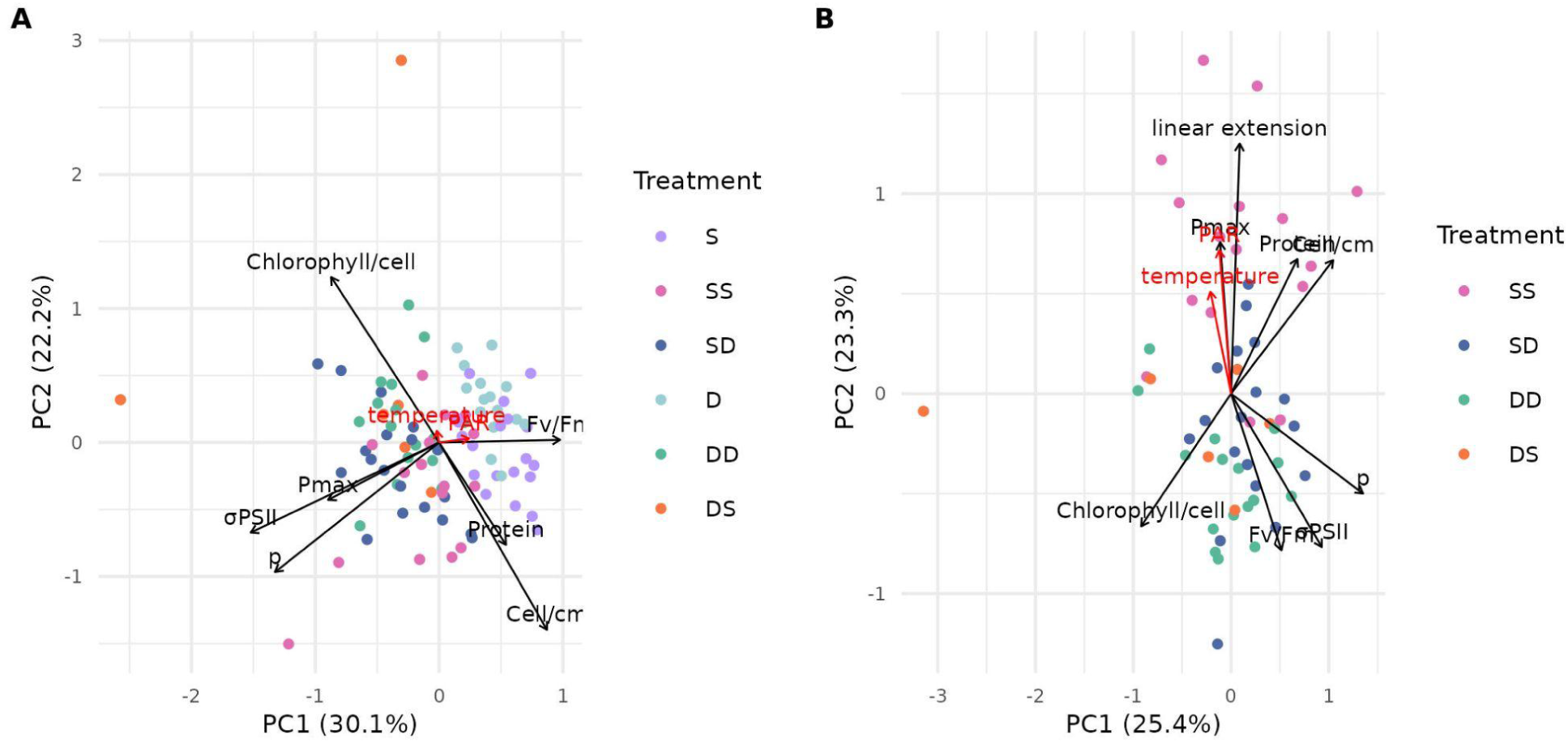
Multivariate structure of skeletal and physiological traits across depth and transplantation groups. **(A)** PCA of physiological variables with fitted environmental vectors measured pre- and post-translocation. **(B)** PCA of physiological and skeletal variables with fitted environmental variables measured post-translocation. Black arrows indicate physiological and skeletal vectors, red arrows indicate fitted environmental vectors. Arrow direction shows the gradient of increasing values and arrow length reflects the strength of correlation with the ordination axes. Variables include functional absorption cross-section (σPSII), connectivity parameter (p), symbiont cells cm^-2^ (Cell/cm), chlorophyll content per symbiont cell (Chlorophyll/cell, µg cell^-1^), photosynthetic efficiency (Fv/Fm), maximum photosynthetic rate (Pmax, electrons s^-1^ PSII^-1^), protein concentration (Protein, µg mL^-1^), skeletal linear extension (cm year^-1^), light availability (PAR, μmol photons m^-2^s^-1^) and temperature (°C).

### Molecular mechanisms underlying coral depth-dependent variation

#### Variation in transcriptomic responses between experimental groups

The molecular basis underlying depth-specific morphology differences in native and translocated corals was assessed using RNA-Seq by testing gene expression in *P. astreoides* samples from various experimental groups. PCA of variance-stabilized (vst) gene counts followed by PERMANOVA showed significant effects of depth and season on overall expression patterns between seasons (Fig. 4A; depth: *P* = 0.001; season: *P* = 0.001). PCA-based transcriptomic shift for SS and DD, calculated as variance-weighted distance from the corresponding winter centroid, did not differ significantly among groups (Fig. 4B).

**Figure 4.**
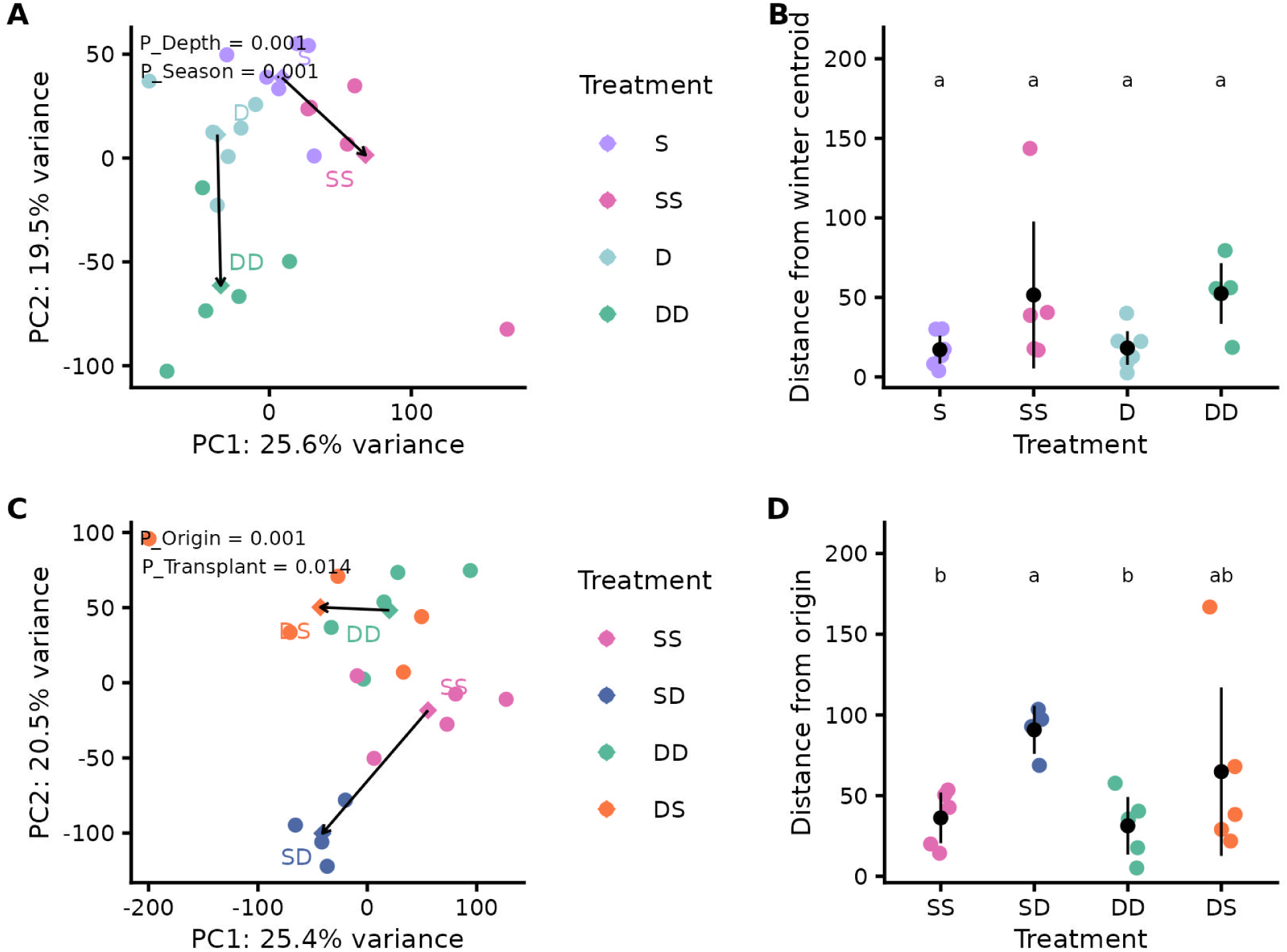
Transcriptomic shifts across seasons and transplantation. **(A)** PCA of vst-counts in winter (S, D) and summer (SS, DD) native sampes. Arrows connect winter S and D centroids to their corresponding summer centroids. Percentages represent the variance explained by each principal component (PC). PERMANOVA results for depth and season are shown. **(B)** Variance-weighted PCA distance from the corresponding winter depth centroid for seasonal comparisons. Points represent individual samples; black points and error bars show group means ± 95% CI. Letters indicate significant differences among groups based on Kruskal-Wallis tests followed by BH-corrected pairwise Wilcoxon tests. **(C)** PCA of vst-counts in summer native (SS, DD) and reciprocal transplant (SD, DS) samples. Arrows connect control centroids to their corresponding transplant centroids. Percentages represent the variance explained by each PC. PERMANOVA results for origin depth and transplant depth are shown. **(D)** Variance-weighted PCA distance from the corresponding origin-depth centroid for transplantation treatments. Points represent individual samples; black points and error bars show group means ± 95% CI. Letters indicate significant differences among groups based on BH-corrected pairwise Wilcoxon tests.

Pairwise differential expression (DE) analysis (p adjusted < 0.05) indicated that depth-associated divergence was limited but greater in summer than in winter, increasing from 0.8% out of 31,989 of expressed genes in the S vs. D contrast to 1.9% in the SS vs. DD contrast (Fig. S8A). To assess the effect of transplantation, PCA followed by PERMANOVA indicated significant effects of origin depth and transplant depth on global expression (Fig. 4C; origin: *P* = 0.001; transplant: *P* = 0.014). SD showed the clearest directional shift from the SS centroid in the PC1–PC2 plane and differed significantly from SS in variance-weighted distance from the origin (Fig. 4D). DS showed an intermediate response: it was not significantly different from DD, but also did not differ from SD, likely because DS corals exhibited great individual-level heterogeneity (Fig. 4D). Venn analyses (Fig. S8B-C) showed that SD vs. DD and SD vs. SS shared a large portion of differentially expressed genes, indicating that a similar set of genes is activated in SD corals regardless of comparison depth.

#### Functional enrichment of differentially expressed genes between experimental groups

Gene Ontology enrichment analysis revealed several GO terms significantly enriched across different contrasts (Fig. 5A, Data S1). Comparisons among S and D in winter showed enrichment of sterol and fatty acid transport processes in shallow corals. In summer, SS corals exhibited upregulation of genes associated with mitochondrial respiration, electron transport, and energy-production pathways compared to DD colonies. DS colonies showed downregulation of structural and extracellular matrix-related tenascin complex genes relative to DD corals as well as reduced expression of genes involved in DNA synthesis and cell-cycle progression compared to SS corals. SD transplants displayed significant downregulation of mRNA degradation and apoptosis-related processes compared to SS corals. GO enrichment performed separately on upregulated and downregulated gene sets recovered the same functional patterns, while identifying more enriched terms within each contrast (Data S2).

**Figure 5.**
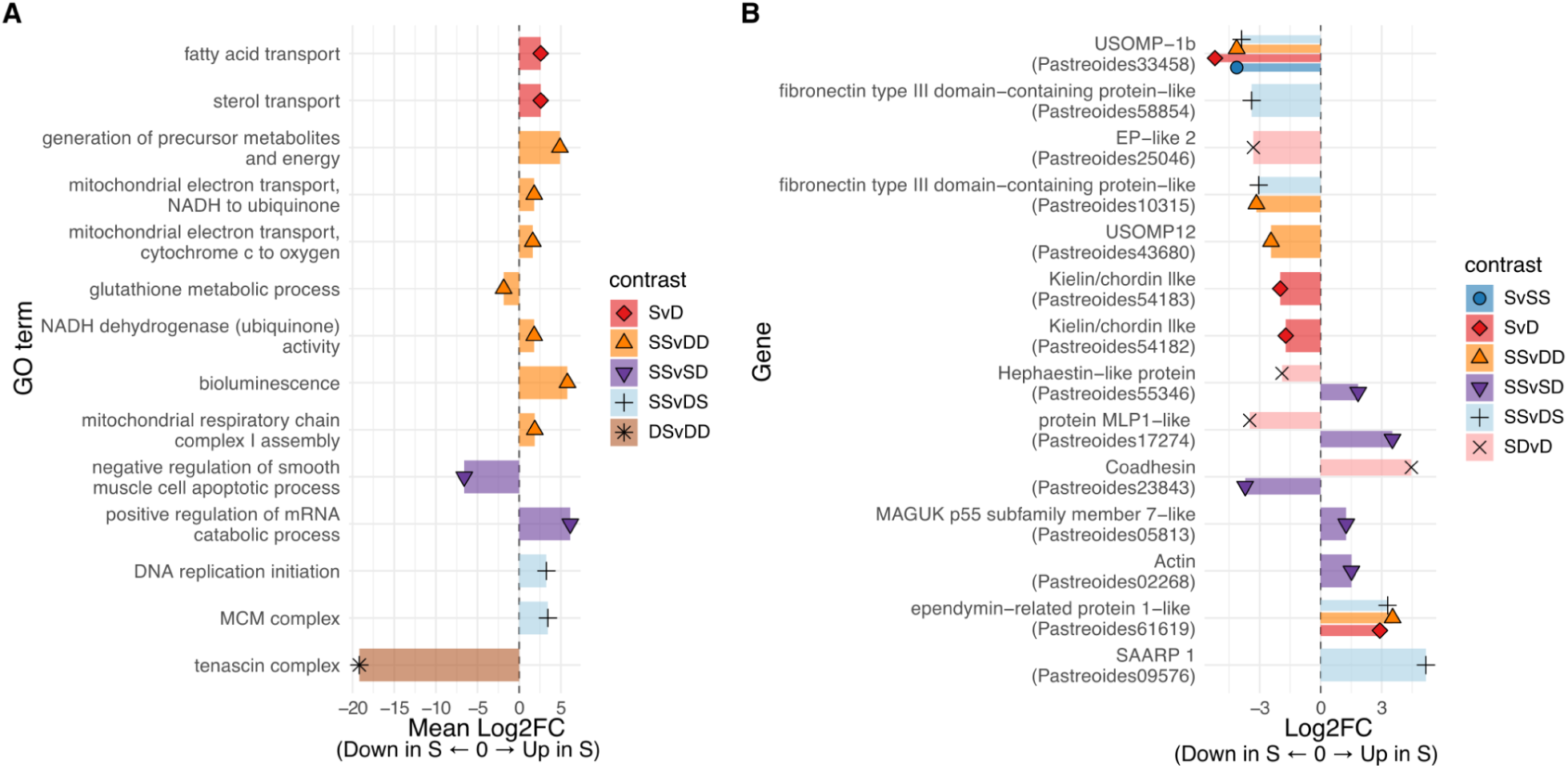
Gene Ontology enrichment and differential expression of biomineralization toolkit genes across experimental groups. **(A)** Gene Ontology (GO) enrichment among DEGs. Bars represent mean log₂ fold change of genes within each enriched term. Non-metazoan GO terms are excluded (see Data S1 for the full table). Positive values indicate higher expression in the first group of each contrast. Contrasts without significantly enriched terms are not shown. **(B)** Differentially expressed biomineralization-related genes. Log_2_ fold change values are shown for coral biomineralization genes identified by orthology. Positive values indicate higher expression in the first group of each contrast. Contrasts without differentially expressed biomineralization genes are not shown.

#### Expression of coral biomineralization-related genes

We evaluated the differential expression of known coral biomineralization-related genes (Fig. 5B, S9). Comparisons between colonies revealed that several structural skeletal organic matrix (SOM) components, including *kielin/chordin-like*, *fibronectin type III domain–containing proteins*, *USOMP-12* (*von Willebrand factor*), and *USOMP-1b*, were consistently downregulated in shallow-origin corals (S, SS, SD) compared to mesophotic groups (D, DD), whereas only an *ependymin-related protein 1-like* was upregulated. The same *ependymin-related protein 1-like* gene remained upregulated in SS corals relative to DS colonies, together with *SAARP1* (skeletal aspartic acid–rich protein 1, also known as *CAPR4*), while *USOMP-1b* and *fibronectin III* remained depleted in SS corals in this comparison. The *MLP1-like viral inclusion protein* gene was strongly depleted in SD corals compared to both SS and DD, mirroring the pattern observed for *hephaestin-like* and opposing that of *coadhesin*, indicating changes associated not to depth but to the treatment itself. The *EP-like 2* gene was enriched in DD compared to SD. Finally, *actin* and *MAGUK p55 subfamily member 7-like* were depleted in SD relative to SS.

#### Co-expression patterns under depth-translocation groups

Broad co-expression patterns associated with depth and translocation treatments were examined using Weighted Correlation Network Analysis (WGCNA). All expressed genes were assigned into 40 co-expression modules, which were further clustered into nine module clusters by eigengene similarity (Fig. 6A). Differences in mean eigengene expression were tested across experimental groups to identify co-expression patterns among clusters (Fig. 6B).

**Figure 6.**
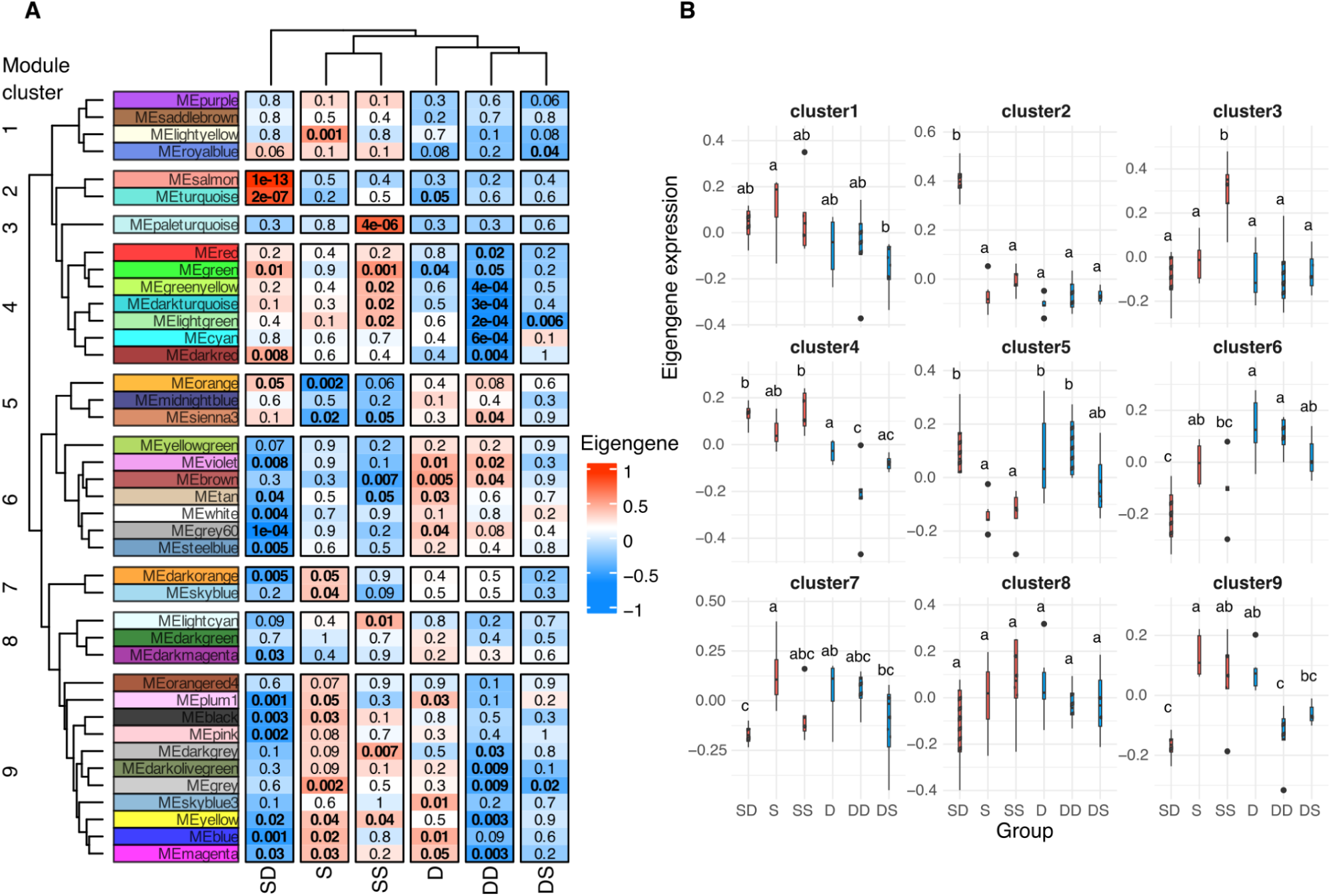
Gene co-expression modules associated with depth and transplantation. **(A)** Heatmap showing correlations between module eigengenes and experimental groups. Values range from −1 (negative correlation) to 1 (positive correlation); significant correlations (p < 0.05) are indicated in bold. Modules are grouped into nine clusters based on eigengene similarity. **(B)** Mean eigengene expression for each module cluster across experimental groups. Points represent individual samples; different letters indicate significant differences (adjusted p < 0.05).

### Functional enrichment of gene co-expression clusters

GO enrichment analysis (Data S3) was performed for genes with significant module-trait correlation within each cluster. Cluster 2, positively correlated to SD transplants, was enriched in DNA repair, recombination, and replication, as well as signaling, transporter activity, and metalloendopeptidase processes. Cluster 3, correlated with SS corals, was enriched for ribosome biogenesis, protein synthesis, folding, and RNA processing. Cluster 4, whose eigengenes were positively correlated with all shallow-origin corals (S, SS, SD), was enriched in mitochondrial oxidative phosphorylation, ATP production, protein translation, proteasome-mediated degradation, transcription, and RNA splicing. Cluster 6, correlated with native deep colonies (D, DD), was enriched for cytoskeletal organization, signaling, transcriptional regulation, and intracellular transport. Cluster 7, associated with S corals, showed enrichment for protein degradation, cellular signaling, and structural remodeling pathways.

## Discussion

Understanding whether vertical persistence in corals is driven primarily by phenotypic plasticity or by fixed depth-specific adaptation is central to predicting reef resilience under climate change. Seasonal fluctuations in photosynthetic performance, energy reserves, and calcification provide a natural context for evaluating coral plasticity (*35*, *36*). In Little Cayman, the study period spanned winter and summer conditions, from November to July, with higher temperatures and greater PAR variability observed during summer (Fig. 1, Table S1–S2). Within-depth seasonal comparisons of molecular and physiological traits in *P. astreoides* showed that significant seasonal variation was mainly restricted to photophysiology, consistent with previous Caribbean observations (*36*). This pattern points to seasonal photoacclimation without broad remodeling within depths, providing a reference for interpreting the reciprocal transplantation experiment.

Depth-associated differences were consistently observed across physiological, skeletal, and transcriptomic traits. Shallow corals exhibited higher metabolic activity, photosynthetic capacity, and linear extension rates, consistent with greater light availability in shallow habitats (*13*, *20*, *21*, *37*). While linear extension reflects one component of the overall calcification process, it does not directly capture changes in skeletal density. In contrast, mesophotic corals displayed reduced tissue protein content and slower skeletal extension, accompanied by elevated expression of skeletal organic matrix (SOM) genes involved in matrix formation. Similar depth-associated shifts in SOM gene expression have been reported in *Stylophora pistillata* (*20*, *25*), suggesting that modulation of skeletal matrix composition represents a conserved strategy under low-light conditions. Upregulation of structural matrix proteins (*38*, *39*) may contribute to maintaining skeletal integrity under reduced mineral deposition, whereas shallow-origin corals showed relatively higher expression of nucleation-associated proteins linked to rapid crystal growth (*40*). These patterns indicate depth-dependent tuning of biomineralization strategies rather than fixed morphological specialization.

Despite pronounced differences in extension rates, internal skeletal thickness was similar across transplantation treatments. This suggests that while linear extension is environmentally plastic, skeletal thickening and density-related traits may represent more conserved structural characteristics (*34*, *41*, *42*). The apparent decoupling between extension and skeletal thickening highlights that calcification encompasses multiple processes, including both volumetric expansion, crystalline deposition and organic matrix scaffolding (*43*, *44*). Such flexibility may allow corals to maintain structural integrity across contrasting light environments, as previously reported (*20*, *25*, *45*). Thus, calcification dynamics appear plastically regulated, while certain aspects of skeletal architecture remain conserved.

Population-level responses to local environmental conditions can arise through genetic adaptation via natural selection and through phenotypic plasticity, the capacity to adjust phenotype in response to environmental variation without changes in genotype (*46*, *47*). In *P. astreoides*, depth-associated genetic differentiation has been documented in the Florida Keys (*48*, *49*), whereas populations in southeast Florida, Bermuda, and the U.S. Virgin Islands exhibit high genetic connectivity across depths (*25*, *48*), indicating that vertical connectivity and the potential for depth-associated genetic divergence vary among and within regions. Cryptic lineages have also been reported in *P. astreoides* and other coral species (*50*), further complicating inference of population structure.

In the Cayman Islands, kinship analysis revealed high clonality and close relatedness within and between depths. High levels of clonality in *P. astreoides* have been reported across multiple locations (*49*, *51*) and are consistent with its capacity for asexual reproduction through fragmentation, self-fertilization, and parthenogenesis (*52–55*). The occurrence of close relatives among colonies randomly sampled from different habitats suggests the presence of genetic exchange between depths and sites. Although SNP-based analyses using large numbers of loci can remain informative under limited sampling (*56*, *57*), the small number of independent genets and the RNA-seq-derived nature of the SNP dataset limit population-level inference about genetic divergence across groups, but does not exclude the possibility of a partial depth-specific specialization.

Connectivity across depths may also be reinforced by symbiont community composition. In both shallow and mesophotic habitats, symbiont community was dominated by *Symbiodinium* (formerly clade A (*58*)), consistent with previous reports from Bermuda (*25*, *48*). This genus exhibits high tolerance to light variability and has been associated with enhanced host performance under fluctuating irradiance (*59*), potentially promoting functional connectivity across depth gradients.

Transplantation responses were strongly asymmetric. SD corals maintained survival rates comparable to SS colonies and exhibited coordinated physiological and transcriptomic shifts toward mesophotic phenotypes, but adjustment to mesophotic conditions involved a distinct remodeling of molecular and skeletal architecture rather than simple adoption of the native morphotype. Activation of apoptosis-regulatory pathways and DNA repair processes (*31*, *60–64*), together with downregulation of catabolic pathways consistent with energy conservation (*65*, *66*), indicates an integrated acclimation program under reduced irradiance. Conversely, DS corals experienced elevated mortality, reduced symbiont density, an indicator of stress and bleaching (*67*), and limited activation of coordinated gene networks. Downregulation of DNA replication–related processes suggests stress-associated cellular disruption rather than adaptive reprogramming (*31*, *68*, *69*). Surviving DS colonies retained mesophotic-like biomineralization expression profiles, suggesting constrained plasticity and indications of depth-specific specialization.

This asymmetry in coral acclimation aligns with theoretical and empirical expectations that exposure to variable environments favors greater phenotypic plasticity (*28–30*). Consistent with the greater environmental heterogeneity in shallow reefs (Fig. 1, Table S1-S2) compared with mesophotic habitats, our findings indicate that phenotypic plasticity decreases with depth, with shallow coral populations exhibiting greater capacity for acclimation to environmental change.

The transcriptional signatures of shallow corals observed here overlap with pathways reported in corals from other environmentally heterogeneous habitats, including shallow, mangrove, high-temperature, turbid, inshore and nearshore reefs (*20*, *25*, *29*, *31*, *32*, *70–74*). Recurrent enrichment of processes related to DNA repair, redox homeostasis, apoptosis regulation, and metabolic remodeling (Fig. 7; Table S7; Data S4) suggests the existence of shared acclimation modules activated under multi-stressor exposure. Corals originating from variable environments may therefore maintain stress-response pathways that can be rapidly activated, allowing them to respond more flexibly to environmental change. The recurrence of these pathways across habitat types suggests that coral resilience relies on common molecular mechanisms rather than habitat-specific responses.

**Figure 7.**
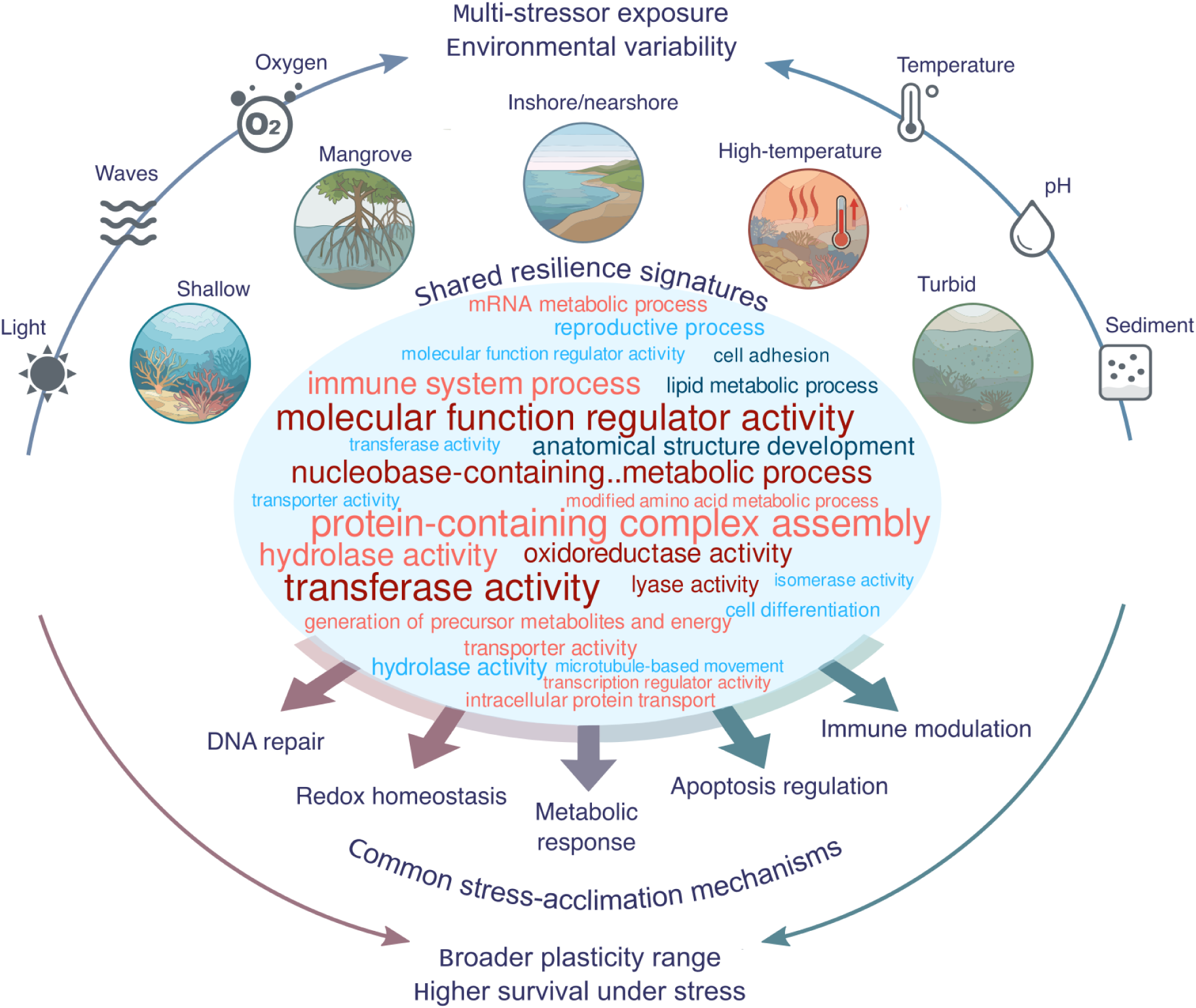
Literature-based synthesis of recurrent functional signatures in corals from environmentally variable habitats. Conceptual model summarizing recurrent functional categories reported across studies of corals inhabiting heterogeneous reef environments exposed to multiple stressors, based on published studies (*20*, *25*, *29*, *31*, *32*, *70–74*). The central word cloud summarizes recurrent GO-slim functional categories identified through differential expression and functional enrichment analyses in the present study and five independent transcriptomic or proteomic studies of corals inhabiting environmentally variable habitats, including shallow reefs (*20*, *25*, *70*), mangrove reefs (*71*), and nearshore reefs (*73*). Word size is proportional to the number of independent studies in which a given term was enriched. Color intensity reflects recurrence consistency across studies. Red shades indicate functional categories upregulated in variable environments; blue shades indicate downregulated categories. Only GO-slim terms detected in at least two independent studies are shown (see Table S7). The full dataset underlying this synthesis is provided in Data S4. For clarity, “nucleobase-containing small molecule metabolic process” is abbreviated in the figure.

These findings refine the Deep Reef Refugia Hypothesis (*10*, *11*). While mesophotic reefs may buffer certain disturbances (*7*, *8*), our results indicate that refugial capacity is conditional on the interplay between phenotypic plasticity and environmental specialization across the depth gradient. Mesophotic-origin *P. astreoides* exhibited constrained upward acclimation, suggesting that deep populations may not universally serve as functional reservoirs for shallow reef recovery under rapid climate change. At the same time, kinship links across depths suggest incomplete genetic isolation, while the greater acclimatory capacity of shallow-origin corals may facilitate their persistence across the depth gradient. Deep-origin coral colonies may require longer and more gradual acclimation in order to succeed (*20*, *75*). Future experiments incorporating stepwise acclimation protocols will help disentangle immediate stress responses from fundamental limits of plasticity.

Collectively, our results suggest that phenotypic plasticity supports vertical persistence in *P. astreoides*, but this plasticity is uneven along the depth gradient. Shallow reefs appear to function as reservoirs of environmentally tolerant phenotypes, whereas mesophotic populations exhibit greater specialization under low-light conditions. As climate change intensifies, reef resilience will depend not only on connectivity across depth gradients but also on how phenotypic plasticity is distributed within and among coral populations.

## Materials and Methods

### Experimental design

The study was conducted on Little Cayman Island, the smallest of the Cayman Islands, with a land area of approximately 25 km^2^, located northwest of Jamaica and south of Cuba in the Caribbean Sea. To increase sample size, two sites with similar geomorphology and temperature conditions, both outside marine protected areas (MPAs), were selected: Martha’s Finyard (MF) (N19.66487, W80.11118) and Coral City (CC) (N19.68075, W80.02330) (Fig. 1A). Both sites comprise sloping inner-reef spur-and-groove formations leading to a near-vertical fore-reef wall extending beyond 2000 m depth. At both sites, temperature and photosynthetic active radiation (PAR; μmol m^-2^ s^-1^) were measured, at 10 and 40 m, using miniPAR loggers equipped with miniWIPER which provided a gentle brush every 6 hours across the PAR logger to eliminate fouling organisms (PME, Vista, California, USA). Temperature was recorded every 30 min, and PAR was recorded every 10 or 15 min. Logger data was available from July to September 2022 for MF site and from April to July 2023 for both sites.

*Porites astreoides*, a brooding Caribbean coral abundant from shallow to mesophotic depths and considered resilient (*76*, *77*), was chosen as a model species to investigate acclimation across the depth gradient. At both locations, 10 colonies of *P. astreoides* were collected at 10 m (S) and 40 m depth (D) on November 27-28, 2022, using a hammer and chisel. Sampled colonies were located at least 5 m from each other to decrease likelihood of sampling clones. Colonies were transferred to the outdoor mesocosm at Central Caribbean Marine Institute (CCMI) where they were sampled and then maintained in flow tanks under ambient temperature and light. All colonies experienced a natural light period. Colonies were cut in half and stained with 1% Alizarin Red for 4-5 hours (*78*). All colonies were then left to recover for 5-6 days before being returned to the reef in a reciprocal transplantation design. Half of each colony was placed back at its native depth (shallow→shallow, SS; deep→deep, DD), while the other half was transplanted to the opposite depth (shallow→deep, SD; deep→shallow, DS). All corals were retrieved on 7-9 July 2023, after ∼7.5 months of the experiment. Dead colonies were excluded from all downstream analyses and were used only for survival estimates.

### Ex situ photophysiological analysis

Ex situ physiological assays were conducted on November 28–29, 2022 (T₀), and again on July 7-9, 2023 (T_end_). After collection, colonies were placed in the mesocosm to recover for 12 h, and active fluorescence was measured in the dark using a diving-fluorescence induction and relaxation (FIRe) fluorometer (*79*). To account for intra-colony variation, five measurements were taken from different positions on each colony and averaged to represent whole-colony photophysiology. The FIRe fluorometer protocol involved photophysiological assessments under both dark-adapted conditions (achieved via shading, irrespective of time of day) and saturating ambient irradiance. Dark-adapted measurements characterize parameters governing photosynthetic performance under light-limiting conditions, specifically the initial slope of the photosynthesis–irradiance curve, whereas assessments under saturating light quantify maximum photosynthetic rates (Pmax, s^-1^ PSII^-1^). This measurements allows for the determination of key photosynthetic parameters, including the maximum quantum yield of PSII photochemistry (Fv/Fm), the functional absorption cross-section of PSII (Sigma: σPSII, in Å²), and the connectivity parameter (ρ), which determines the probability of excitation energy transfer between adjacent PSII units (dimensionless) (*80*).

### Ex situ physiological analyses

After collecting, a piece of about 5 cm was cut from each coral, snap-frozen and kept in −20 °C for analysis of symbiont density, chlorophyll concentration, and protein content. Each fragment was airbrushed into a sterile ziplock bag containing 4 mL of distilled water, and the resulting tissue slurry was transferred into 15-mL centrifuge tubes and electrically homogenized for 20 s. Homogenates were centrifuged at 5,000 × g for 10 min at 4 °C, and a 100-µL aliquot of the supernatant was used to quantify host protein concentration using the QPRO-BCA kit (Cyanagen, Italy) according to the manufacturer’s instructions. Total protein concentration was measured at 562 nm using a PerkinElmer 2300 EnSpire plate reader. Following an additional centrifugation step (5,000 × g for 5 min at 4°C), removal of the supernatant, and resuspension of the pellet in filtered seawater, 50-µL aliquots were used to determine algal symbiont density. Symbiont counts were obtained by fluorescence microscopy with a hemocytometer (n = 4 per sample) and imaged in both brightfield and under 440-nm excitation (Nikon Eclipse Ti–S Inverted Microscope System) to visualize chlorophyll and ensure that only symbiont cells were counted. Surface area was measured using ImageJ for the fragment image. Resulting cell densities were normalized to surface area for each fragment. Chlorophyll a concentrations were quantified from 2 mL of homogenate incubated overnight in 90% cold acetone at 4 °C. Samples were then centrifuged at 5,000 × g for 5 min at 4°C, and absorbance was measured in a 96-well microplate. Chlorophyll concentrations were calculated using the equations of Jeffrey and Humphrey (*81*), with path length adjusted to 0.555 cm for a 200-µL well volume.

### Skeletal micromorphology

#### Skeletal extension rate analysis and skeletal morphology

Skeletal extension rates were quantified using Alizarin Red, a fluorescent dye widely employed to mark coral skeletal growth (*78*, *82*). During calcification, the dye becomes incorporated into the calcium carbonate skeleton, forming a distinct band that serves as a temporal marker.

The cleaned skeletons remaining after the physiology assay described above were imaged using a Nikon stereomicroscope (SMZ800N), and skeletal extension was measured as the linear distance between the upper edge of the septa and the bottom of the alizarin line. For each coral fragment, this distance was measured three times, and the mean value was used as the estimate of skeletal growth over the 7.5-month period. In addition, each fragment was imaged using Leica Emspira 3 microscope (Leica, Microsystems, Wetzlar, Germany). For each fragment corallite diameter and corallite density were measured.

#### Micro–Computed Tomography (Micro-CT) Imaging

Coral skeletons were imaged using laboratory micro–computed tomography (micro-CT) instruments (EasyTom S, RX Solutions, France) to obtain three-dimensional reconstructions of skeletal architecture. Scans were acquired at a 15-µm pixel size, over 360°, with frame averaging of 5 and a frame rate of 4 s^-1^. The beam was set to 100 kV, 296 uA, and filtered with 0.3 mm Cu. Datasets were reconstructed using X-Act (RX Solutions, France) with a fixed dynamic range, and ring filtering was applied with a kernel of 5. Reconstructed datasets were examined in 2D and 3D using ImageJ (National Institutes of Health, USA) to verify structural completeness and orientation before analysis. Image stacks were imported into Fiji (ImageJ), resliced and rotated to align the skeletal growth axis vertically. A volume of interest of ∼313 mm^3^ was cropped to include both older (prior to alizarin staining) and new skeletal material. Segmentation of skeletal material and void space was performed using the Trainable Weka Segmentation plugin (*83*), for which approximately 20 representative slices were manually annotated to define “skeleton” and “void” classes before training and applying the classifier to the full dataset. The resulting segmented stacks were converted to binary format (skeleton = 255; void = 0), referred to as coral volume masks. Skeleton thickness was quantified from these masks using the BoneJ Slice Geometry module in Fiji, with voxel dimensions confirmed from acquisition metadata; thickness was calculated as Mean Thickness 3D (µm) for each slice. The porosity of each volume of interest was calculated as the percent of pore volume relative to total volume, using a custom python script (https://doi.org/10.5281/zenodo.22080924). Total volume masks were created by filling all voids within the coral volume masks, both open and closed to the outside (2D morphological closing, radius = 20 pixels). Pore masks were then derived as the ‘difference’ (logical and-not) between the total volume mask and the coral volume mask. Finally, porosity was calculated by counting the pixels included (labelled 255) in the pore and total volume masks and taking their ratio.

### RNA extraction and sequencing

Tissue samples of 1 cm^2^ were taken from each coral fragment before the translocation (groups S and D) and in the end point (groups SS, SD, DD and DS) and preserved in DNA/RNA Shield (Zymo Research). Samples were stored at −80 °C until processing. Total RNA was extracted using a Bio-TriRNA (Molecular Biology) with an addition of 2-mercaptoethanol (Aldrich) and chloroform combined with Quick-DNA/RNA Miniprep Plus Kit (Zymo Research) spin-column purification. Coral samples were gently scratched with a sterile razor blade and subsequently treated with Bio-TriRNA, 2-mercaptoethanol and chloroform. After vortexing and centrifugation, the clear upper phase of RNA was transferred to a green Zymo spin-column, purified and DNAse-treated following the manufacturer’s protocol. At the final elution step, 30 μl of RNAse-free water was passed twice through the spin column to maximize the concentration of eluted RNA. RNA concentration was measured using a NanoDrop 2000 (Thermo Fisher Scientific) and RNA integrity was tested on a TapeStation (Agilent Technologies). Illumina library preparation and sequencing were conducted by GENEWIZ, LLC (Azenta Life Sciences). Paired-end 150 bp reads were generated using an Illumina NovaSeq 6000 platform. In total, 31 samples were sequenced with a median of 33.5 million reads per sample. The resulting sample group sizes were: S (n=6), D (n=6), SS (n=5), SD (n=4), DD (n=5), DS (n=5). Raw reads for all samples are available on the NCBI Sequence Read Archive (SRA) under BioProject number PRJNA1426050.

### Bioinformatic analysis

#### RNA-seq filtering, mapping and symbiont species identification

Raw read quality was assessed using FastQC v0.12.1 (*84*) and summarized with MultiQC v1.0 (*85*) (Table S8). Reads were trimmed with Trimmomatic v0.39 (*86*) to remove adapters and low-quality bases and retain reads > 25 bp in length. rRNA percentage was evaluated using SortMeRNA v4.3.6 (*87*). Filtered reads were aligned to the *Porites astreoides* genome assembly (SRR19144705) (*88*) using STAR v2.7.10b (*89*) with *--quantMode GeneCounts* option to obtain gene-level read counts. The absence of a 3’ bias was validated with RSEQC v5.0.4 (*90*). Across samples, 1.4–21.1 million reads mapped uniquely to the *P. astreoides* genome (median = 7.3 million), of which 0.5–12.1 million reads were assigned to genes (median = 3.8 million). For the taxonomic screening of a non-host read fraction, unmapped holobiont reads were subsequently aligned to a proteome database of Symbiodiniaceae species (Table S9) using DIAMOND v0.9 (*91*) and e-value threshold of 0.01. For each read, only the best hit was considered. The proportion of reads mapped to each symbiont proteome was then calculated, visualized in R v4.4.2 (*92*) and additionally summarized across symbiont genera.

#### Host variants discovery and SNP-based analyses

Single nucleotide polymorphisms (SNPs) were identified from the host reads using the Genome Analysis Toolkit (GATK v4.6.2) (*93*), following the Broad Institute’s best practices for RNA-seq variant discovery (*94*), with modifications for non-model organisms lacking known variant databases. Reads aligned to *P. astreoides* genome were sorted and marked for duplicates prior to variant calling with GATK HaplotypeCaller. Joint genotyping was then performed using GenotypeGVCFs, and variant sets were filtered for quality by depth using SelectVariants and VariantFiltration, yielding approximately 69,000 high-confidence SNPs. SNPs were pruned for linkage disequilibrium in PLINK v2.0 (*95*). Fragments originating from the same colony (i.e. artificial replicates) were collapsed to a single representative to retain only independent biological samples, and variants were filtered by a minor allele frequency ≥0.05 and a maximum missing rate of 0.4, resulting in a dataset of 6,337 SNPs across 20 individuals (S.MF: n=5, S.CC: n=5, D.MF: n=4, D.CC: n=6). Genetic relatedness among colonies was evaluated using KING kinship coefficients calculated in PLINK. Clonemates were defined using a pairwise kinship threshold > 0.354, and first-degree relatives were identified using a threshold > 0.177. The filtered genotype matrix was converted into a Genomic Data Structure (GDS) format in R using the SNPRelate package v1.42.0 (*96*). Related samples were collapsed to retain one representative per genet, resulting in a clone/relative-filtered dataset of 13 individuals (S.CC: n=4, D.MF: n=4, D.CC: n=5). Pairwise fixation indices (Fst, (*97*)) were calculated using StAMPP v1.6.3 (*98*) with 1,000 bootstrap replicates.

### PCA-based analysis of seasonal and transplantation-induced transcriptomic shifts

Transcriptomic shifts were assessed using a PCA-based distance analysis following (*99*). Two analyses were performed separately. First, seasonal transcriptomic shifts were assessed using winter (S, D) and summer native (SS, DD) samples. Second, transplantation-induced shifts were assessed using summer native (SS, DD) and transplant samples (SD, DS). Low-count raw reads were filtered to keep genes with >=10 counts in at least five (seasonal) or four (translocation) samples. Filtered count matrices were normalized using a variance-stabilizing transformation (vst) in DESeq2 v1.44 (*100*), and effects of site, rRNA content, and RIN score were removed using limma::removeBatchEffect. PCA was performed on the corrected vst-counts. Permutational multivariate analysis of variance (PERMANOVA) was used to test whether gene expression patterns were influenced by depth, season, and their interaction in the seasonal analysis, and by origin depth, transplant depth, and their interaction in the transplantation analysis. Transcriptomic shifts were quantified across the first two PCs as variance-weighted Euclidean distances to each sample from the corresponding origin centroid: winter S or D centroids for the seasonal dataset, and summer SS or DD centroids for the transplantation dataset. Distance metrics were compared among groups using Kruskal-Wallis tests followed by Benjamini-Hochberg-corrected pairwise Wilcoxon tests.

#### Differential expression analysis and biomineralization gene profiling

Analyses were carried out in the R statistical environment. Low-expression genes (those with less than 10 counts in less than four samples) were removed from the dataset, leaving 31,989 *P. astreoides* genes. Gene counts were normalized and log-transformed with *rlog* command in DESeq2 v1.44. The effect of the experimental group was analyzed with an account to the site and to the quality of RNA sequences (*design = ∼ site + rRNA + RIN + condition*). Because paired samples were retained for only a subset of colonies, transcriptomic analysis was performed using an unpaired design after confirming that within-colony Bray-Curtis distances did not differ significantly from between-colony distances. Differentially expressed genes were identified in each contrast using Wald test with an adjusted p-value (padj) < 0.05. Venn diagrams were plotted with the ggVennDiagram v1.5.0 R package (*101*). The *P. astreoides* orthologs of the known coral biomineralization-related genes (*38–40*, *102–104*) were identified using Proteinortho v6.2.3 (*105*) with an e-value cutoff of 1×10⁻¹⁰, yielding 106 genes associated with biomineralization, and compared with the differential expression results. Significantly differentially expressed genes among these were detected for each contrast.

#### Weighted Gene Co-expression Network Analysis

For the Weighted Gene Co-expression Network Analysis (WGCNA) (*106*), filtered gene counts were normalized using the variance stabilizing transformation (vst) in DESeq2. Because WGCNA is sensitive to outliers, samples were hierarchically clustered using the R function *hclust*, and two outlier samples were removed, leaving 29 samples in the analysis: S (n=6), D (n=6), SS (n=5), SD (n=4), DD (n=5), DS (n=3). A soft threshold power of 11 was selected using *pickSoftThreshold* in the WGCNA R package v1.73 and used for the blockwiseModules automatic network construction function with minModuleSize = 30, adjacency = “signed” and mergeCutHeight = 0.2, resulting in 40 co-expression modules. Modules were further clustered by eigengene similarity into nine module clusters using hierarchical clustering. Expression profiles for each cluster were summarized as mean eigengene expression per experimental group and visualized using ComplexHeatmap v2.24 (*107*). Variation in eigengene expression between experimental groups was assessed with limma v3.64 (*108*) with Benjamini–Hochberg multiple correction (p.adjusted < 0.05). Module-trait correlations were calculated by comparing gene significance (correlation between genes and experimental group) with module membership (correlation between module eigengenes and gene expression). Genes within each module were considered significant if |module membership| > 0.8 (p < 0.05) and gene significance > 0.2 (p < 0.05).

#### Gene ontology enrichment analysis

Gene Ontology (GO) enrichment analysis was performed with the clusterProfiler v4.16.0 R package (*109*) to identify over-representation of particular GO terms in both DE contrasts and WGCNA module clusters. Since *P. astreoides* is a non-model species, custom TERM2GENE and TERM2NAME tables were generated using the available *P. astreoides* GO annotation (*108*) and the go.obo database (released on 16.03.2025) (*110*). All expressed *P. astreoides* genes served as a background. Over-representation analysis was carried out using the *enricher* function (pvalueCutoff = 0.05, pAdjustMethod = “fdr”, qvalueCutoff = 0.2) for three gene set types: (i) significantly differentially expressed genes identified by DESeq2 for each contrast (adjusted p value < 0.05 and |log₂FoldChange| > 1), (ii) direction-specific subsets of significant DE genes analyzed separately for upregulated and downregulated genes within each contrast, and (iii) significant genes from each WGCNA module cluster. GO terms with an adjusted p value (FDR) < 0.05 were considered significantly enriched. Mean log₂ fold change values were calculated for genes comprising each term from set (i) and (ii).

To identify recurrent transcriptomic signatures across studies, GO enrichment results from the present study and five additional transcriptomic and proteomic studies (*20*, *25*, *70*, *71*, *73*) comparing corals from environmentally variable and stable habitats were compiled into a unified dataset and mapped to GO slim categories. For each study, only GO terms reported as significantly enriched according to the criteria defined in the original publication were retained. Terms enriched in corals from more variable habitats (e.g., shallow, inshore, or mangrove lagoon) compared to corals from more stable environments (e.g., mesophotic, offshore, or reef) were classified as upregulated, whereas the reverse contrast was classified as downregulated. GO slim terms were then aggregated across studies, retaining only categories detected in at least two independent studies. For each term, total occurrences across studies were calculated to determine word size, and the number of studies in which each term appeared was used to define color intensity. Terms were separated by direction of enrichment.

### Statistical analysis

Survival was analyzed at the individual level using a binomial generalized linear model with treatment and site as additive fixed effects, followed by Tukey-adjusted post hoc comparisons. Physiological, photophysiological and skeletal parameters were tested for normality (Shapiro-Wilk test) and homogeneity of variances (Levene’s test). Where necessary, data were root-square or log-transformed. For variables meeting parametric assumptions, linear mixed-effects models (LMM) were used. When assumptions were not met, alternative approaches were used: an inverse Gaussian GLMM for chl/cell; LMM with fitted group-specific residual variance (*weights = varIdent(form = ∼ 1 | Group)*, *method = "REML"*) for σPSII, connectivity (p), Fv/Fm and Pmax (Table S3). Tukey-adjusted post hoc comparisons were performed for all models. In every analysis, group was included as a fixed effect, and colony identity as a random effect. Corallite diameter was analyzed using an LMM with sample as a random effect, since multiple corallites were measured per fragment. Corallite density (number of polyps per 5 mm²) was analyzed using a Poisson GLM. Physiological parameters, transformed when necessary, were used as an input to a PCA. Environmental parameters (temperature and PAR) were subsequently fitted as vectors onto the ordination space. Group differences in physiological profiles between experimental groups were tested with PERMANOVA with Euclidean distances calculated from transformed and scaled data and 999 permutations. For samples after transplantation, PERMANOVA was performed using an origin × transplant depth design. Pairwise comparisons between experimental groups were further assessed using pairwise PERMANOVA. To compare Symbiodiniaceae community composition between coral groups, Bray-Curtis dissimilarities were calculated based on relative abundance profiles at both the species and genus levels, and group differences were tested with PERMANOVA (10000 permutations). A Wilcoxon rank-sum test was applied to compare within- and between-group compositional dissimilarity. All differences were considered significant at adjusted p < 0.05. All computations were performed using R v4.5.1. The R and Python codes and raw data are available on github https://github.com/talimass/Cayman-translocation and citable at Zenodo https://doi.org/10.5281/zenodo.22080924.

## Acknowledgments

We thank Dr. Rachel Bober (University of Haifa) for her expert guidance and assistance with RNA extraction, and Dr. Maya Lalzar (University of Haifa) for her consultations on statistical analysis. We are grateful to Jeanne Bloomberg and Kimberly Malkoski for their assistance during fieldwork. ES acknowledges support from the Center for Integration in Science, Ministry of Aliyah and Integration, and a postdoctoral scholarship from the University of Haifa.

## Author contributions

Conceptualization: GG-G, HN, SE, TM

Data curation: ES, GG-G, SE, PZ, AC

Formal analysis: ES, IV, PZ, TM

Funding acquisition: GG-G, TM

Investigation: ES, GG-G, HN, SE, PZ, AC, TM

Methodology: ES, GG-G, HN, SE, IV, PZ, AC, TM

Project administration: GG-G, AC, TM

Resources: GG-G, IV, AC, TM

Software: ES, IV, TM

Supervision: GG-G, AC, TM

Validation: GG-G, TM

Visualization: ES, HN, IV, PZ, TM

Writing – original draft: ES, TM

Writing – review & editing: ES, GG-G, SE, IV, PZ, AC, TM

## Data, code and materials availability

All data needed to evaluate the conclusions in the paper are present in the paper, the Supplementary Materials and online at https://doi.org/10.5281/zenodo.22080924. Raw sequencing data have been deposited in the NCBI SRA under BioProject accession PRJNA1426050. R code for all analyses presented in this study can be accessed at https://doi.org/10.5281/zenodo.22080924. This study did not generate new materials.

## References

1. R. W. Grigg, J. J. Polovina, M. J. Atkinson, Model of a coral reef ecosystem. Coral Reefs 3, 23–27 (1984).

2. R. Wood, Reef Evolution (Oxford University Press, 1999).

3. T. A. Gardner, I. M. Côté, J. A. Gill, A. Grant, A. R. Watkinson, Long-term region-wide declines in Caribbean corals. Science 301, 958–960 (2003).

4. G. De’ath, K. E. Fabricius, H. Sweatman, M. Puotinen, The 27-year decline of coral cover on the Great Barrier Reef and its causes. Proc Natl Acad Sci U S A 109, 17995–17999 (2012).

5. G. J. Edgar, R. D. Stuart-Smith, F. J. Heather, N. S. Barrett, E. Turak, H. Sweatman, M. J. Emslie, D. J. Brock, J. Hicks, B. French, S. C. Baker, S. A. Howe, A. Jordan, N. A. Knott, P. Mooney, A. T. Cooper, E. S. Oh, G. A. Soler, C. Mellin, S. D. Ling, J. C. Dunic, J. W. Turnbull, P. B. Day, M. F. Larkin, Y. Seroussi, J. Stuart-Smith, E. Clausius, T. R. Davis, J. Shields, D. Shields, O. J. Johnson, Y. H. Fuchs, L. Denis-Roy, T. Jones, A. E. Bates, Continent-wide declines in shallow reef life over a decade of ocean warming. Nature 615, 858–865 (2023).

6. C. Cai, N. M. Hammerman, J. M. Pandolfi, C. M. Duarte, S. Agusti, Influence of global warming and industrialization on coral reefs: A 600-year record of elemental changes in the Eastern Red Sea. Sci Total Environ 914, 169984 (2024).

7. L. A. Rocha, H. T. Pinheiro, B. Shepherd, Y. P. Papastamatiou, O. J. Luiz, R. L. Pyle, P. Bongaerts, Mesophotic coral ecosystems are threatened and ecologically distinct from shallow water reefs. Science 361, 281–284 (2018).

8. C. Diaz, N. L. Foster, M. J. Attrill, A. Bolton, P. Ganderton, K. L. Howell, E. Robinson, P. Hosegood, Mesophotic coral bleaching associated with changes in thermocline depth. Nat Commun 14, 6528 (2023).

9. L. M. Hinderstein, J. C. A. Marr, F. A. Martinez, M. J. Dowgiallo, K. A. Puglise, R. L. Pyle, D. G. Zawada, R. Appeldoorn, Theme section on “Mesophotic Coral Ecosystems: Characterization, Ecology, and Management.” Coral Reefs 29, 247–251 (2010).

10. P. Bongaerts, T. Ridgway, E. M. Sampayo, O. Hoegh-Guldberg, Assessing the ‘deep reef refugia’ hypothesis: focus on Caribbean reefs. Coral Reefs 29, 309–327 (2010).

11. M. P. Lesser, M. Slattery, C. D. Mobley, Biodiversity and Functional Ecology of Mesophotic Coral Reefs. Annual Review of Ecology, Evolution, and Systematics 49, 49–71 (2018).

12. R. K. Cowen, S. Sponaugle, Larval dispersal and marine population connectivity. Ann Rev Mar Sci 1, 443–466 (2009).

13. T. Mass, S. Einbinder, E. Brokovich, N. Shashar, R. Vago, J. Erez, Z. Dubinsky, Photoacclimation of Stylophora pistillata to light extremes: metabolism and calcification. Marine Ecology Progress Series 334, 93–102 (2007).

14. G. Eyal, J. Wiedenmann, M. Grinblat, C. D’Angelo, E. Kramarsky-Winter, T. Treibitz, O. Ben-Zvi, Y. Shaked, T. B. Smith, S. Harii, V. Denis, T. Noyes, R. Tamir, Y. Loya, Spectral Diversity and Regulation of Coral Fluorescence in a Mesophotic Reef Habitat in the Red Sea. PLOS ONE 10, e0128697 (2015).

15. M. P. Lesser, M. Slattery, J. J. Leichter, Ecology of mesophotic coral reefs. Journal of Experimental Marine Biology and Ecology 375, 1–8 (2009).

16. J. F. Bruno, P. J. Edmunds, Clonal Variation for Phenotypic Plasticity in the Coral Madracis Mirabilis. Ecology 78, 2177–2190 (1997).

17. S. Einbinder, T. Mass, E. Brokovich, Z. Dubinsky, J. Erez, D. Tchernov, Changes in morphology and diet of the coral Stylophora pistillata along a depth gradient. Marine Ecology Progress Series 381, 167–174 (2009).

18. G. Goodbody-Gringley, C. Marchini, A. D. Chequer, S. Goffredo, Population Structure of Montastraea cavernosa on Shallow versus Mesophotic Reefs in Bermuda. PLoS One 10, e0142427 (2015).

19. G. Goodbody-Gringley, J. Waletich, Morphological plasticity of the depth generalist coral, Montastraea cavernosa, on mesophotic reefs in Bermuda. Ecology 99, 1688–1690 (2018).

20. A. Malik, S. Einbinder, S. Martinez, D. Tchernov, S. Haviv, R. Almuly, P. Zaslansky, I. Polishchuk, B. Pokroy, J. Stolarski, T. Mass, Molecular and skeletal fingerprints of scleractinian coral biomineralization: From the sea surface to mesophotic depths. Acta Biomaterialia 120, 263–276 (2021).

21. M. L. Doherty, A. D. Chequer, T. Mass, G. Goodbody-Gringley, Phenotypic variability of Montastraea cavernosa and Porites astreoides along a depth gradient from shallow to mesophotic reefs in the Cayman Islands. Coral Reefs 43, 1173–1187 (2024).

22. T. J. DeWitt, S. M. Scheiner, Eds., Phenotypic Plasticity: Functional and Conceptual Approaches (Oxford University Press, 2004; 10.1093/oso/9780195138962.001.0001).

23. A. R. A. Mateus, M. Marques-Pita, V. Oostra, E. Lafuente, P. M. Brakefield, B. J. Zwaan, P. Beldade, Adaptive developmental plasticity: Compartmentalized responses to environmental cues and to corresponding internal signals provide phenotypic flexibility. BMC Biol 12, 97 (2014).

24. G. E. Carpenter, A. D. Chequer, S. Weber, T. Mass, G. Goodbody-Gringley, Light and photoacclimatization drive distinct differences between shallow and mesophotic coral communities. Ecosphere 13, e4200 (2022).

25. F. Scucchia, K. Wong, P. Zaslansky, H. M. Putnam, G. Goodbody-Gringley, T. Mass, Morphological and genetic mechanisms underlying the plasticity of the coral *Porites astreoides* across depths in Bermuda. Journal of Structural Biology 215, 108036 (2023).

26. L. Eyal-Shaham, G. Eyal, R. Tamir, Y. Loya, Reproduction, abundance and survivorship of two Alveopora spp. in the mesophotic reefs of Eilat, Red Sea. Sci Rep 6, 20964 (2016).

27. X. Pochon, Z. H. Forsman, H. L. Spalding, J. L. Padilla-Gamiño, C. M. Smith, R. D. Gates, Depth specialization in mesophotic corals (Leptoseris spp.) and associated algal symbionts in Hawai’i. R Soc Open Sci. 2, 140351 (2015).

28. T. J. Dewitt, A. Sih, D. S. Wilson, Costs and limits of phenotypic plasticity. Trends Ecol Evol 13, 77–81 (1998).

29. C. D. Kenkel, M. V. Matz, Gene expression plasticity as a mechanism of coral adaptation to a variable environment. Nat Ecol Evol 1, 14 (2016).

30. L.-M. Chevin, A. A. Hoffmann, Evolution of phenotypic plasticity in extreme environments. Philos Trans R Soc Lond B Biol Sci 372, 20160138 (2017).

31. D. J. Barshis, J. T. Ladner, T. A. Oliver, F. O. Seneca, N. Traylor-Knowles, S. R. Palumbi, Genomic basis for coral resilience to climate change. Proceedings of the National Academy of Sciences 110, 1387–1392 (2013).

32. R. A. Bay, S. R. Palumbi, Transcriptome predictors of coral survival and growth in a highly variable environment. Ecol Evol 7, 4794–4803 (2017).

33. W. Johnston, A. Cooper, Small islands and climate change: analysis of adaptation policy in the Cayman Islands. Reg Environ Change 22, 45 (2022).

34. P. S. Davies, Effect of daylight variations on the energy budgets of shallow-water corals. Mar. Biol. 108, 137–144 (1991).

35. W. K. Fitt, F. K. McFarland, M. E. Warner, G. C. Chilcoat, Seasonal patterns of tissue biomass and densities of symbiotic dinoflagellates in reef corals and relation to coral bleaching. Limnology and Oceanography 45, 677–685 (2000).

36. L. Chapron, V. Schoepf, S. J. Levas, M. D. Aschaffenburg, M. E. Warner, A. G. Grottoli, Natural Variability in Caribbean Coral Physiology and Implications for Coral Bleaching Resilience. Front. Mar. Sci. 8 (2022).

37. M. P. Lesser, M. Slattery, M. Stat, M. Ojimi, R. D. Gates, A. Grottoli, Photoacclimatization by the coral Montastraea cavernosa in the mesophotic zone: light, food, and genetics. Ecology 91, 990–1003 (2010).

38. J. L. Drake, T. Mass, L. Haramaty, E. Zelzion, D. Bhattacharya, P. G. Falkowski, Proteomic analysis of skeletal organic matrix from the stony coral Stylophora pistillata. Proceedings of the National Academy of Sciences 110, 3788–3793 (2013).

39. M. P. Mummadisetti, J. L. Drake, P. G. Falkowski, The spatial network of skeletal proteins in a stony coral. J R Soc Interface 18, 20200859 (2021).

40. T. Mass, J. L. Drake, L. Haramaty, J. D. Kim, E. Zelzion, D. Bhattacharya, P. G. Falkowski, Cloning and Characterization of Four Novel Coral Acid-Rich Proteins that Precipitate Carbonates In Vitro. Current Biology 23, 1126–1131 (2013).

41. T. M. DeCarlo, A. L. Cohen, Dissepiments, density bands and signatures of thermal stress in Porites skeletons. Coral Reefs 36, 749–761 (2017).

42. D. J. Barnes, J. M. Lough, On the nature and causes of density banding in massive coral skeletons. Journal of Experimental Marine Biology and Ecology 167, 91–108 (1993).

43. F. Scucchia, P. Harnay, H. M. Putnam, Contrasting temperature-induced gene network rewiring and isoform switching underlie relative thermal tolerance of coral species. bioRxiv [Preprint] (2025). 10.64898/2025.12.26.696607.

44. P. U. P. A. Gilbert, K. D. Bergmann, N. Boekelheide, S. Tambutté, T. Mass, F. Marin, J. F. Adkins, J. Erez, B. Gilbert, V. Knutson, M. Cantine, J. O. Hernández, A. H. Knoll, Biomineralization: Integrating mechanism and evolutionary history. Sci Adv 8, eabl9653 (2022).

45. F. Scucchia, A. Malik, P. Zaslansky, H. M. Putnam, T. Mass, Combined responses of primary coral polyps and their algal endosymbionts to decreasing seawater pH. Proc Biol Sci 288, 20210328 (2021).

46. E. Sanford, M. W. Kelly, Local Adaptation in Marine Invertebrates. Annual Review of Marine Science 3, 509–535 (2011).

47. O. Savolainen, M. Lascoux, J. Merilä, Ecological genomics of local adaptation. Nat Rev Genet 14, 807–820 (2013).

48. X. M. Serrano, I. B. Baums, T. B. Smith, R. J. Jones, T. L. Shearer, A. C. Baker, Long distance dispersal and vertical gene flow in the Caribbean brooding coral Porites astreoides. Sci Rep 6, 21619 (2016).

49. E. N. Shilling, R. J. Eckert, A. B. Sturm, J. D. Voss, Porites astreoides coral populations demonstrate high clonality and connectivity in southeast Florida. Coral Reefs 42, 1131–1145 (2023).

50. G. N. Cabacungan, T. N. W. Kankanamalage, A. F. Azam, M. R. Collins, A. R. Arratia, A. N. Gutting, M. V. Matz, K. L. Black, Cryptic coral community composition across environmental gradients. PLOS ONE 20, e0318653 (2025).

51. F. Riquet, A. Japaud, F. L. D. Nunes, X. M. Serrano, A. C. Baker, E. Bezault, C. Bouchon, C. Fauvelot, Complex spatial patterns of genetic differentiation in the Caribbean mustard hill coral Porites astreoides. Coral Reefs 41, 813–828 (2022).

52. D. A. Brazeau, D. F. Gleason, M. E. Morgan, Self-fertilization in brooding hermaphroditic Caribbean corals: Evidence from molecular markers. Journal of Experimental Marine Biology and Ecology 231, 225–238 (1998).

53. D. F. Gleason, D. A. Brazeau, D. Munfus, Can self-fertilizing coral species be used to enhance restoration of Caribbean reefs? Bulletin of Marine Science 69, 933–943 (2001).

54. G. Goodbody-Gringley, S. De Putron, “Brooding Corals: Planulation Patterns, Larval Behavior, and Recruitment Dynamics in the Face of Environmental Change” (2016), pp. 279–289.

55. A. Vollmer, Rare Parthenogenic Reproduction in a Common Reef Coral, Porites astreoides. HCNSO Student Theses and Dissertations (2018).

56. E.-M. Willing, C. Dreyer, C. van Oosterhout, Estimates of Genetic Differentiation Measured by FST Do Not Necessarily Require Large Sample Sizes When Using Many SNP Markers. PLOS ONE 7, e42649 (2012).

57. A. G. Nazareno, J. B. Bemmels, C. W. Dick, L. G. Lohmann, Minimum sample sizes for population genomics: an empirical study from an Amazonian plant species. Mol Ecol Resour 17, 1136–1147 (2017).

58. T. C. LaJeunesse, J. E. Parkinson, P. W. Gabrielson, H. J. Jeong, J. D. Reimer, C. R. Voolstra, S. R. Santos, Systematic Revision of Symbiodiniaceae Highlights the Antiquity and Diversity of Coral Endosymbionts. Current Biology 28, 2570–2580.e6 (2018).

59. J. M. Reynolds, B. U. Bruns, W. K. Fitt, G. W. Schmidt, Enhanced photoprotection pathways in symbiotic dinoflagellates of shallow-water corals and other cnidarians. Proc Natl Acad Sci U S A 105, 13674–13678 (2008).

60. D. Tchernov, H. Kvitt, L. Haramaty, T. S. Bibby, M. Y. Gorbunov, H. Rosenfeld, P. G. Falkowski, Apoptosis and the selective survival of host animals following thermal bleaching in zooxanthellate corals. Proceedings of the National Academy of Sciences 108, 9905–9909 (2011).

61. H. Kvitt, H. Rosenfeld, K. Zandbank, D. Tchernov, Regulation of Apoptotic Pathways by Stylophora pistillata (Anthozoa, Pocilloporidae) to Survive Thermal Stress and Bleaching. PLOS ONE 6, e28665 (2011).

62. A. Moya, L. Huisman, S. Forêt, J.-P. Gattuso, D. C. Hayward, E. E. Ball, D. J. Miller, Rapid acclimation of juvenile corals to CO2-mediated acidification by upregulation of heat shock protein and Bcl-2 genes. Molecular Ecology 24, 438–452 (2015).

63. M. Guo, T. Yuan, L. Jiang, G. Zhou, H. Huang, Acclimation mechanisms of reef-building coral Acropora gemmifera juveniles to long-term CO2-driven ocean acidification. Sci Rep 15, 30655 (2025).

64. C. D. Roper, D. J. Suggett, K. Songsomboon, J. Edmondson, H. England, T. D. Haydon, S. Goyen, C. M. Duijser, R. Alderdice, C. R. Voolstra, E. F. Camp, Coral thermotolerance retained following year-long exposure to a novel environment. Science Advances 11, eadu3858 (2025).

65. I. M. Sokolova, M. Frederich, R. Bagwe, G. Lannig, A. A. Sukhotin, Energy homeostasis as an integrative tool for assessing limits of environmental stress tolerance in aquatic invertebrates. Marine Environmental Research 79, 1–15 (2012).

66. J. Vidal-Dupiol, N. M. Dheilly, R. Rondon, C. Grunau, C. Cosseau, K. M. Smith, M. Freitag, M. Adjeroud, G. Mitta, Thermal Stress Triggers Broad Pocillopora damicornis Transcriptomic Remodeling, while Vibrio coralliilyticus Infection Induces a More Targeted Immuno-Suppression Response. PLOS ONE 9, e107672 (2014).

67. B. E. Brown, Coral bleaching: causes and consequences. Coral Reefs 16, S129–S138 (1997).

68. O. L. Kantidze, A. K. Velichko, A. V. Luzhin, S. V. Razin, Heat Stress-Induced DNA Damage. Acta Naturae 8, 75–78 (2016).

69. I. Yuyama, T. Higuchi, T. Mezaki, H. Tashiro, K. Ikeo, Metatranscriptomic analysis of corals inoculated with tolerant and non-tolerant symbiont exposed to high temperature and light stress. Frontiers in physiology 13, 806171 (2022).

70. M. S. Studivan, J. D. Voss, Transcriptomic plasticity of mesophotic corals among natural populations and transplants of Montastraea cavernosa in the Gulf of Mexico and Belize. Molecular Ecology 29, 2399–2415 (2020).

71. F. Scucchia, P. Zaslansky, C. Boote, A. Doheny, T. Mass, E. F. Camp, The role and risks of selective adaptation in extreme coral habitats. Nat Commun 14, 4475 (2023).

72. K. D. Castillo, C. B. Bove, A. M. Hughes, M. E. Powell, J. B. Ries, S. W. Davies, Gene expression plasticity facilitates acclimatization of a long-lived Caribbean coral across divergent reef environments. Sci Rep 14, 7859 (2024).

73. K. H. Tisthammer, E. Timmins-Schiffman, F. O. Seneca, B. L. Nunn, R. H. Richmond, Physiological and molecular responses of lobe coral indicate nearshore adaptations to anthropogenic stressors. Sci Rep 11, 3423 (2021).

74. E. Bollati, Y. Rosenberg, N. Simon-Blecher, R. Tamir, O. Levy, D. Huang, Untangling the molecular basis of coral response to sedimentation. Molecular Ecology 31, 884–901 (2022).

75. I. Cohen, Z. Dubinsky, Long term photoacclimation responses of the coral Stylophora pistillata to reciprocal deep to shallow transplantation: photosynthesis and calcification. Front. Mar. Sci. 2 (2015).

76. D. Green, P. Edmunds, R. Carpenter, Increasing relative abundance of Porites astreoides on Caribbean reefs mediated by an overall decline in coral cover. Mar. Ecol. Prog. Ser. 359, 1–10 (2008).

77. L. F. O. Lima, H. Bursch, E. A. Dinsdale, Win some, lose some: The ecophysiology of Porites astreoides as a key coral species to Caribbean reefs. Front. Mar. Sci. 9 (2022).

78. Lamberts, A. E., “Coral growth: alizarin method” in Coral Reefs: Research Methods (UNESCO, Paris, 1978), pp. 523–527.

79. Gorbunov M.Y., Falkowski P.G., “Fluorescence Induction and Relaxation (FIRe) Technique and Instrumentation for Monitoring Photosynthetic Processes and Primary Production in Aquatic Ecosystems.” in “Photosynthesis: Fundamental Aspects to Global Perspectives” *(Eds:* A. van Der Est and D. Bruce) (Allen Press), pp. 1029–1031.

80. M. Y. Gorbunov, P. G. Falkowski, Using chlorophyll fluorescence kinetics to determine photosynthesis in aquatic ecosystems. Limnology and Oceanography 66, 1–13 (2021).

81. S. W. Jeffrey, G. F. Humphrey, New spectrophotometric equations for determining chlorophylls *a*, *b*, *c*1 and *c*2 in higher plants, algae and natural phytoplankton. Biochemie und Physiologie der Pflanzen 167, 191–194 (1975).

82. M. Holcomb, A. L. Cohen, D. C. McCorkle, An evaluation of staining techniques for marking daily growth in scleractinian corals. Journal of Experimental Marine Biology and Ecology 440, 126–131 (2013).

83. I. Arganda-Carreras, V. Kaynig, C. Rueden, K. W. Eliceiri, J. Schindelin, A. Cardona, H. Sebastian Seung, Trainable Weka Segmentation: a machine learning tool for microscopy pixel classification. Bioinformatics 33, 2424–2426 (2017).

84. S. Andrews, FastQC: a quality control tool for high throughput sequence data. (2010).

85. P. Ewels, M. Magnusson, S. Lundin, M. Käller, MultiQC: Summarize analysis results for multiple tools and samples in a single report. Bioinformatics, doi: 10.1093/bioinformatics/btw354 (2016).

86. A. M. Bolger, M. Lohse, B. Usadel, Trimmomatic: A flexible trimmer for Illumina sequence data. Bioinformatics 30, 2114–2120 (2014).

87. E. Kopylova, L. Noé, H. Touzet, SortMeRNA: fast and accurate filtering of ribosomal RNAs in metatranscriptomic data. Bioinformatics 28, 3211–3217 (2012).

88. K. H. Wong, H. M. Putnam, The genome of the mustard hill coral, Porites astreoides. GigaByte 2022, gigabyte65 (2022).

89. A. Dobin, C. A. Davis, F. Schlesinger, J. Drenkow, C. Zaleski, S. Jha, P. Batut, M. Chaisson, T. R. Gingeras, STAR: Ultrafast universal RNA-seq aligner. Bioinformatics 29, 15–21 (2013).

90. L. Wang, S. Wang, W. Li, RSeQC: quality control of RNA-seq experiments. Bioinformatics 28, 2184–2185 (2012).

91. B. Buchfink, C. Xie, D. H. Huson, Fast and sensitive protein alignment using DIAMOND. Nat Methods 12, 59–60 (2015).

92. R Core Team, R: A Language and Environment for Statistical Computing. R Foundation for Statistical Computing, (2025); https://www.R-project.org/.

93. A. McKenna, M. Hanna, E. Banks, A. Sivachenko, K. Cibulskis, A. Kernytsky, K. Garimella, D. Altshuler, S. Gabriel, M. Daly, M. A. DePristo, The Genome Analysis Toolkit: a MapReduce framework for analyzing next-generation DNA sequencing data. Genome Res 20, 1297–1303 (2010).

94. G. A. Van der Auwera, M. O. Carneiro, C. Hartl, R. Poplin, G. Del Angel, A. Levy-Moonshine, T. Jordan, K. Shakir, D. Roazen, J. Thibault, E. Banks, K. V. Garimella, D. Altshuler, S. Gabriel, M. A. DePristo, From FastQ data to high confidence variant calls: the Genome Analysis Toolkit best practices pipeline. Curr Protoc Bioinformatics 43, 11.10.1–11.10.33 (2013).

95. S. Purcell, B. Neale, K. Todd-Brown, L. Thomas, M. A. R. Ferreira, D. Bender, J. Maller, P. Sklar, P. I. W. de Bakker, M. J. Daly, P. C. Sham, PLINK: a tool set for whole-genome association and population-based linkage analyses. Am J Hum Genet 81, 559–575 (2007).

96. X. Zheng, D. Levine, J. Shen, S. M. Gogarten, C. Laurie, B. S. Weir, A high-performance computing toolset for relatedness and principal component analysis of SNP data. Bioinformatics 28, 3326–3328 (2012).

97. B. S. Weir, C. C. Cockerham, ESTIMATING F-STATISTICS FOR THE ANALYSIS OF POPULATION STRUCTURE. Evolution 38, 1358–1370 (1984).

98. L. W. Pembleton, N. O. I. Cogan, J. W. Forster, StAMPP: an R package for calculation of genetic differentiation and structure of mixed-ploidy level populations. Molecular Ecology Resources 13, 946–952 (2013).

99. C. B. Bove, K. Greene, S. Sugierski, N. G. Kriefall, A. K. Huzar, A. M. Hughes, K. Sharp, N. D. Fogarty, S. W. Davies, Exposure to global change and microplastics elicits an immune response in an endangered coral. Front. Mar. Sci. 9 (2023).

100. M. I. Love, W. Huber, S. Anders, Moderated estimation of fold change and dispersion for RNA-seq data with DESeq2. Genome Biology 15, 550 (2014).

101. C.-H. Gao, C. Chen, T. Akyol, A. Dusa, G. Yu, B. Cao, P. Cai, ggVennDiagram: Intuitive Venn diagram software extended. iMeta 3, e177 (2024).

102. P. Ramos-Silva, J. Kaandorp, L. Huisman, B. Marie, I. Zanella-Cléon, N. Guichard, D. J. Miller, F. Marin, The Skeletal Proteome of the Coral Acropora millepora: The Evolution of Calcification by Co-Option and Domain Shuffling. Molecular Biology and Evolution 30, 2099–2112 (2013).

103. T. Takeuchi, L. Yamada, C. Shinzato, H. Sawada, N. Satoh, Stepwise Evolution of Coral Biomineralization Revealed with Genome-Wide Proteomics and Transcriptomics. PLOS ONE 11, e0156424 (2016).

104. Y. Peled, J. L. Drake, A. Malik, R. Almuly, M. Lalzar, D. Morgenstern, T. Mass, Optimization of skeletal protein preparation for LC–MS/MS sequencing yields additional coral skeletal proteins in Stylophora pistillata. BMC Mat 2, 8 (2020).

105. M. Lechner, S. Findeiß, L. Steiner, M. Marz, P. F. Stadler, S. J. Prohaska, Proteinortho: Detection of (Co-)orthologs in large-scale analysis. BMC Bioinformatics 12, 1–9 (2011).

106. P. Langfelder, S. Horvath, WGCNA: an R package for weighted correlation network analysis. BMC Bioinformatics 9, 559 (2008).

107. Z. Gu, R. Eils, M. Schlesner, Complex heatmaps reveal patterns and correlations in multidimensional genomic data. Bioinformatics 32, 2847–2849 (2016).

108. M. E. Ritchie, B. Phipson, D. Wu, Y. Hu, C. W. Law, W. Shi, G. K. Smyth, limma powers differential expression analyses for RNA-sequencing and microarray studies. Nucleic Acids Res 43, e47 (2015).

109. G. Yu, L.-G. Wang, Y. Han, Q.-Y. He, clusterProfiler: an R Package for Comparing Biological Themes Among Gene Clusters. OMICS 16, 284–287 (2012).

110. S. A. Aleksander, J. Balhoff, S. Carbon, J. M. Cherry, H. J. Drabkin, D. Ebert, M. Feuermann, P. Gaudet, N. L. Harris, D. P. Hill, R. Lee, H. Mi, S. Moxon, C. J. Mungall, A. Muruganugan, T. Mushayahama, P. W. Sternberg, P. D. Thomas, K. Van Auken, J. Ramsey, D. A. Siegele, R. L. Chisholm, P. Fey, M. C. Aspromonte, M. V. Nugnes, F. Quaglia, S. Tosatto, M. Giglio, S. Nadendla, G. Antonazzo, H. Attrill, G. dos Santos, S. Marygold, V. Strelets, C. J. Tabone, J. Thurmond, P. Zhou, S. H. Ahmed, P. Asanitthong, D. Luna Buitrago, M. N. Erdol, M. C. Gage, M. Ali Kadhum, K. Y. C. Li, M. Long, A. Michalak, A. Pesala, A. Pritazahra, S. C. C. Saverimuttu, R. Su, K. E. Thurlow, R. C. Lovering, C. Logie, S. Oliferenko, J. Blake, K. Christie, L. Corbani, M. E. Dolan, H. J. Drabkin, D. P. Hill, L. Ni, D. Sitnikov, C. Smith, A. Cuzick, J. Seager, L. Cooper, J. Elser, P. Jaiswal, P. Gupta, P. Jaiswal, S. Naithani, M. Lera-Ramirez, K. Rutherford, V. Wood, J. L. De Pons, M. R. Dwinell, G. T. Hayman, M. L. Kaldunski, A. E. Kwitek, S. J. F. Laulederkind, M. A. Tutaj, M. Vedi, S.-J. Wang, P. D’Eustachio, L. Aimo, K. Axelsen, A. Bridge, N. Hyka-Nouspikel, A. Morgat, S. A. Aleksander, J. M. Cherry, S. R. Engel, K. Karra, S. R. Miyasato, R. S. Nash, M. S. Skrzypek, S. Weng, E. D. Wong, E. Bakker, T. Z. Berardini, L. Reiser, A. Auchincloss, K. Axelsen, G. Argoud-Puy, M.-C. Blatter, E. Boutet, L. Breuza, A. Bridge, C. Casals-Casas, E. Coudert, A. Estreicher, M. Livia Famiglietti, M. Feuermann, A. Gos, N. Gruaz-Gumowski, C. Hulo, N. Hyka-Nouspikel, F. Jungo, P. Le Mercier, D. Lieberherr, P. Masson, A. Morgat, I. Pedruzzi, L. Pourcel, S. Poux, C. Rivoire, S. Sundaram, A. Bateman, E. Bowler-Barnett, H. Bye-A-Jee, P. Denny, A. Ignatchenko, R. Ishtiaq, A. Lock, Y. Lussi, M. Magrane, M. J. Martin, S. Orchard, P. Raposo, E. Speretta, N. Tyagi, K. Warner, R. Zaru, A. D. Diehl, R. Lee, J. Chan, S. Diamantakis, D. Raciti, M. Zarowiecki, M. Fisher, C. James-Zorn, V. Ponferrada, A. Zorn, S. Ramachandran, L. Ruzicka, M. Westerfield, The Gene Ontology knowledgebase in 2023. Genetics 224, iyad031 (2023).

