## Supplementary material for "Asymmetric Depth Acclimation and Plasticity Limit the Refugial Potential of Mesophotic *Porites astreoides*": Figs. S1 to S9; Tables S1 to S9

Elizaveta Skalon *et al.*

*

**This PDF file includes:**

Figs. S1 to S9

Tables S1 to S9

**Other Supplementary Materials for this manuscript include the following:**

Data S1 to S4

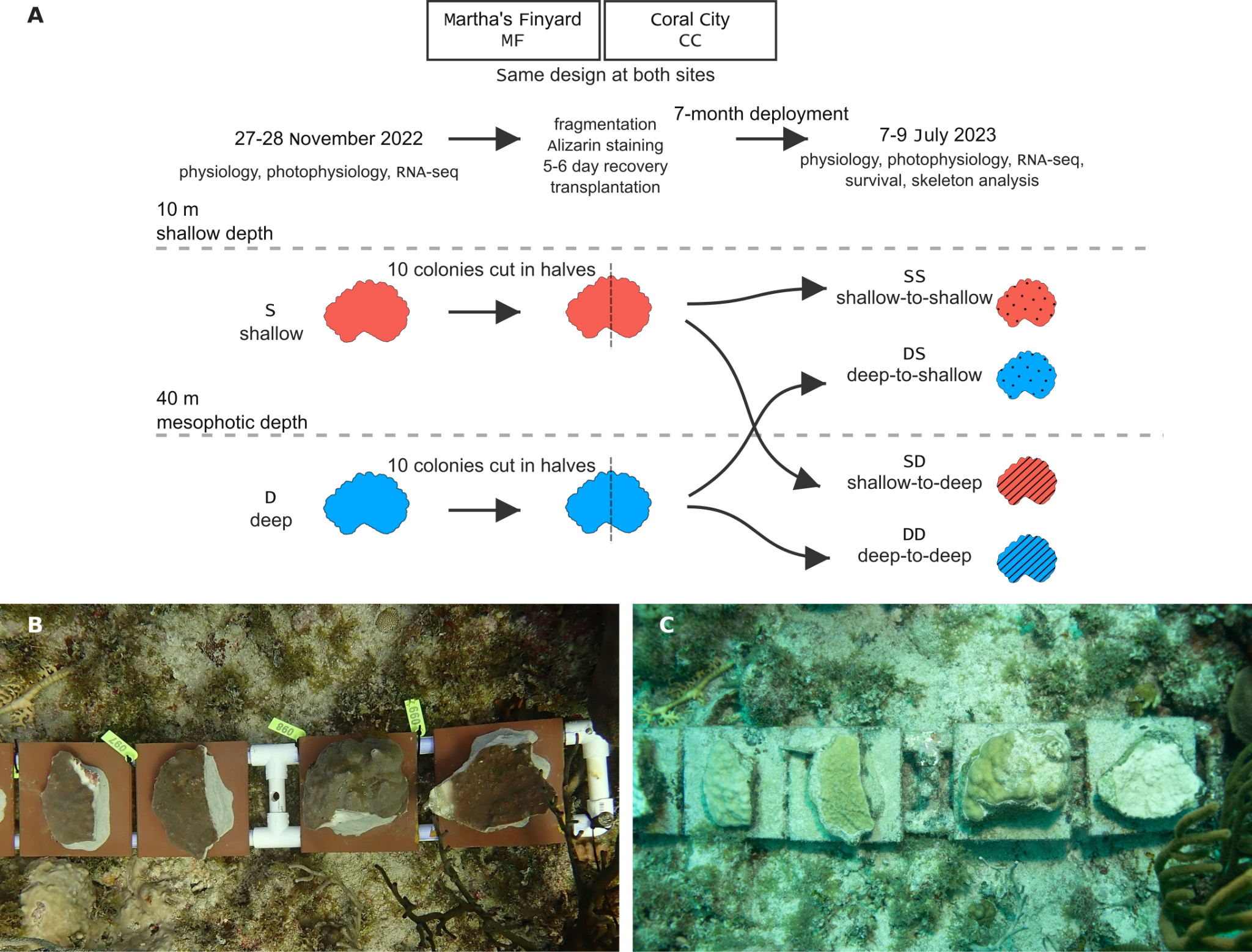

**Fig. S1.** Experimental design and transplanted coral colonies. (A) Schematic representation of the experimental design. (B) Transplanted colonies at the start of the experiment. (C) Transplanted colonies at the end of the experiment.

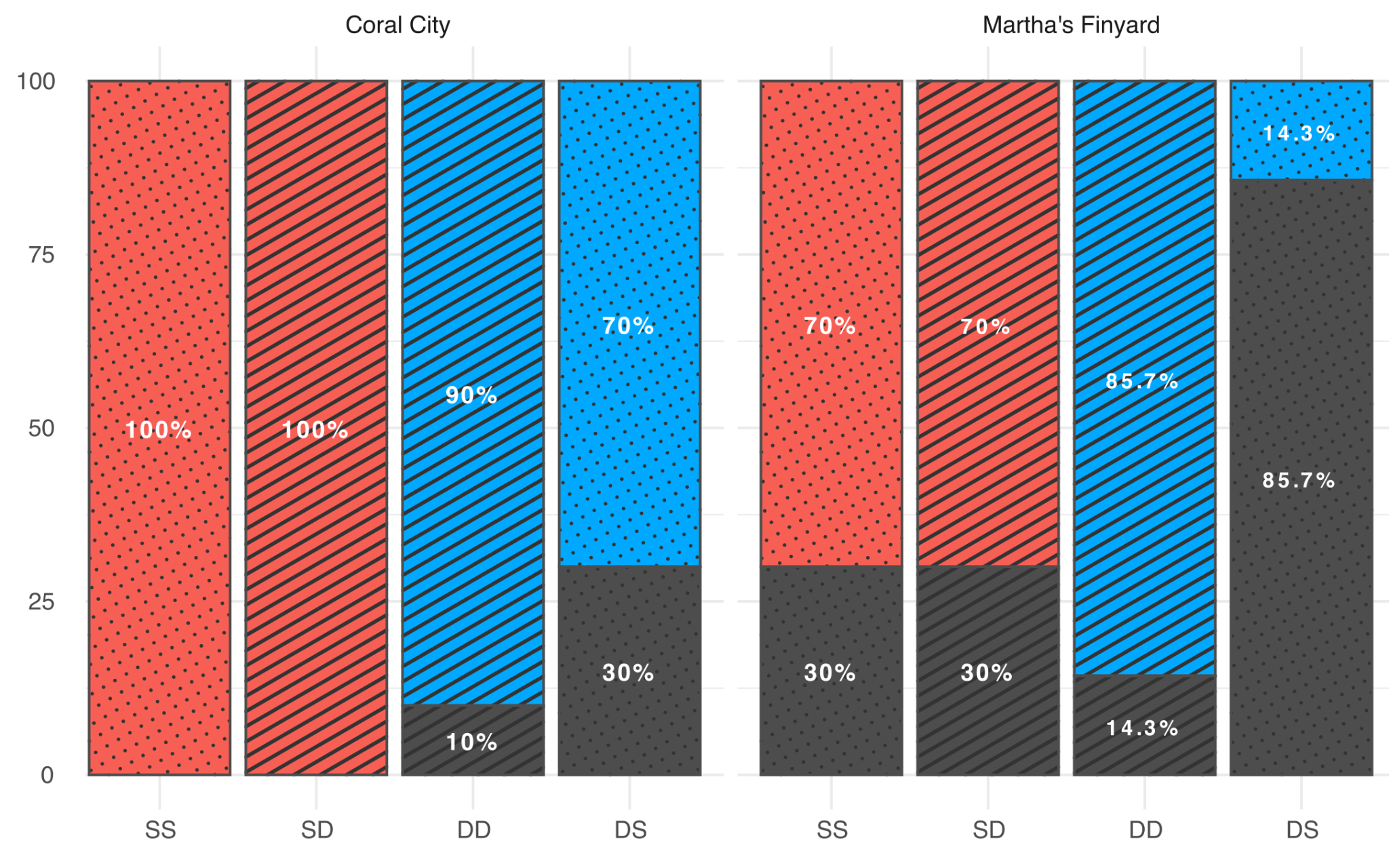

**Fig. S2.** Survival outcomes (alive vs. dead) of *Porites astreoides* colonies across four translocation treatments at two sites (Coral City and Martha’s Finyard). Bars show the percentages of survived (colored) and dead (gray) colonies within each treatment. A site-adjusted logistic regression indicated significant effects of both treatment (χ² = 13.59, p = 0.0035) and site (χ² = 11.99, p = 0.00053) on mortality, with DS transplants exhibiting significantly higher mortality than SS, SD, and DD treatments (Tukey-adjusted p < 0.05).

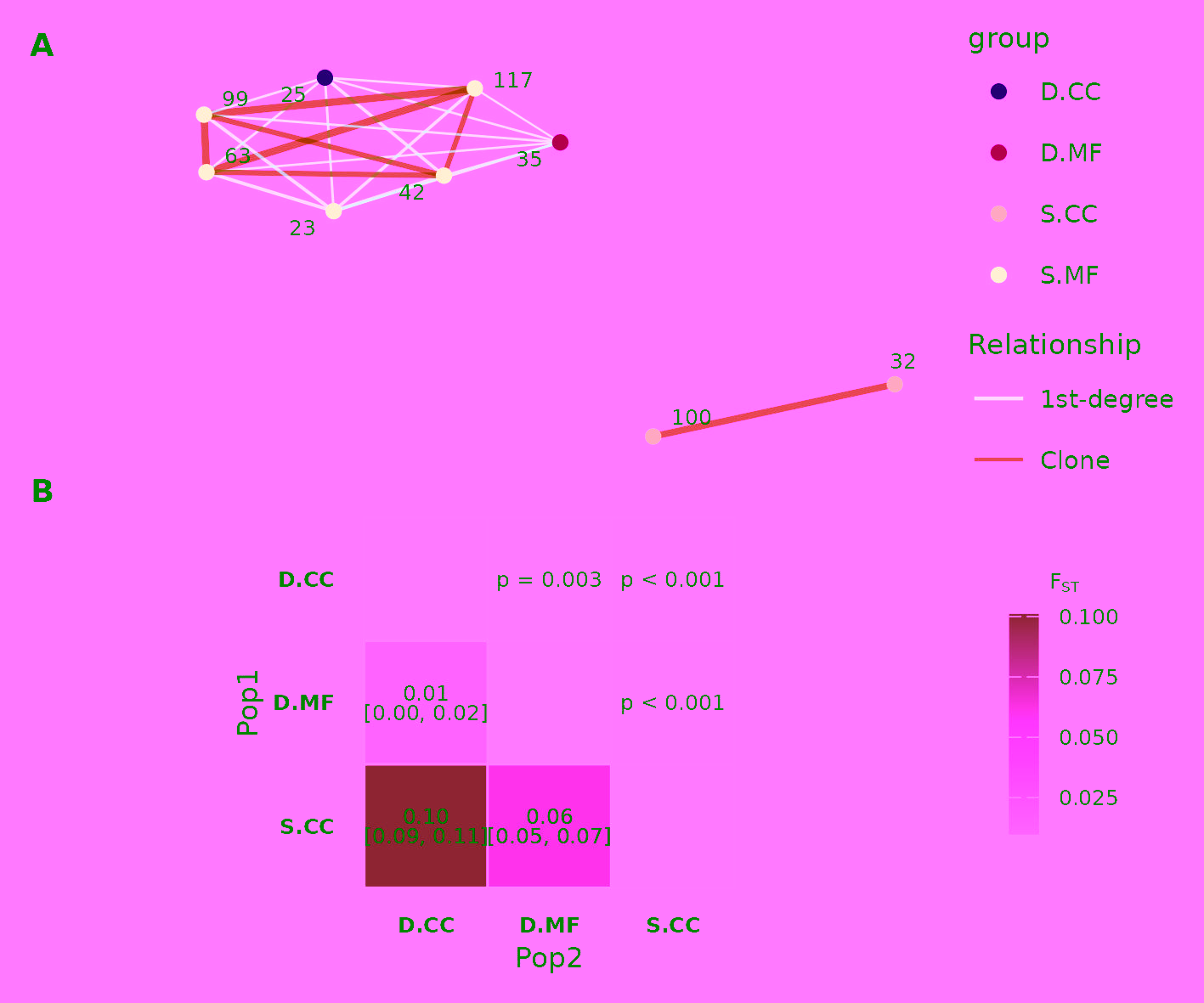

**Fig. S3. Genetic relatedness among *Porites astreoides* colonies across depths and sites. (A)** KING kinship network showing inferred clonal and first-degree relationships. Artificial clones (i.e. fragments of the same colony) are collapsed. Nodes represent individual genets; edges indicate KING kinship coefficients. **(B)** Pairwise Fst estimates calculated after collapsing clonemates and first-degree relatives identified by KING. Values below the diagonal show pairwise Fst with 95% bootstrap confidence intervals; values above the diagonal show p-values based on 1000 bootstrap replicates. The S.MF group was excluded from pairwise Fst analysis because clone/relative filtering reduced its effective sample size. Sample sizes: S.CC: n=4, D.MF: n=4, D.CC: n=5.

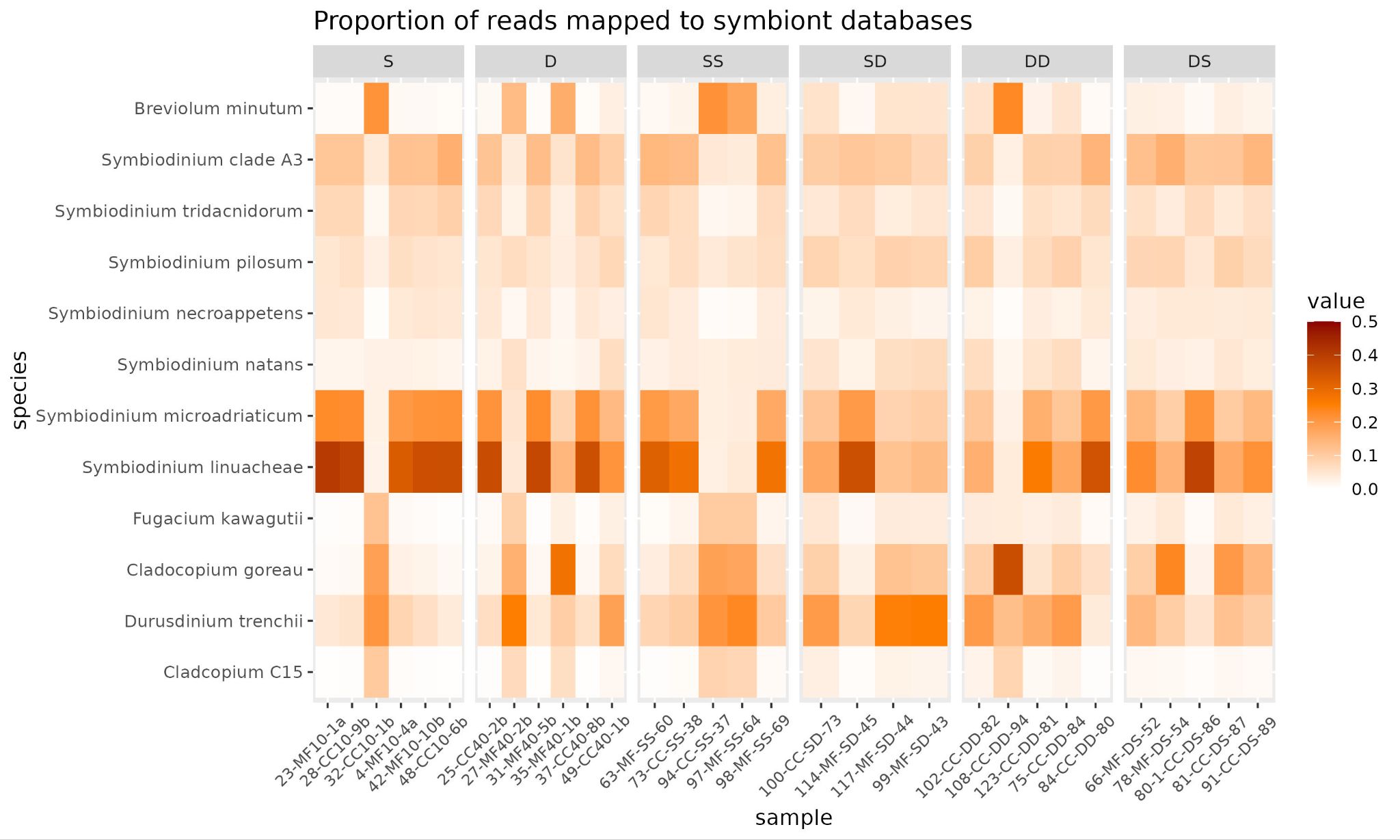

**Fig. S4.** Symbiodiniaceae community composition across treatments. Heatmap showing the proportion of sequencing reads mapped to reference databases of Symbiodiniaceae taxa across individual coral samples. Rows represent symbiont species and columns represent individual samples combined according to the experimental group. Color intensity indicates the relative proportion of reads assigned to each taxon within a sample. Multivariate analysis based on Bray–Curtis dissimilarities detected no significant differences in symbiont community composition among treatments, sites, or depths (PERMANOVA, R² = 0.23, F = 0.59, p = 0.853; 10,000 permutations).

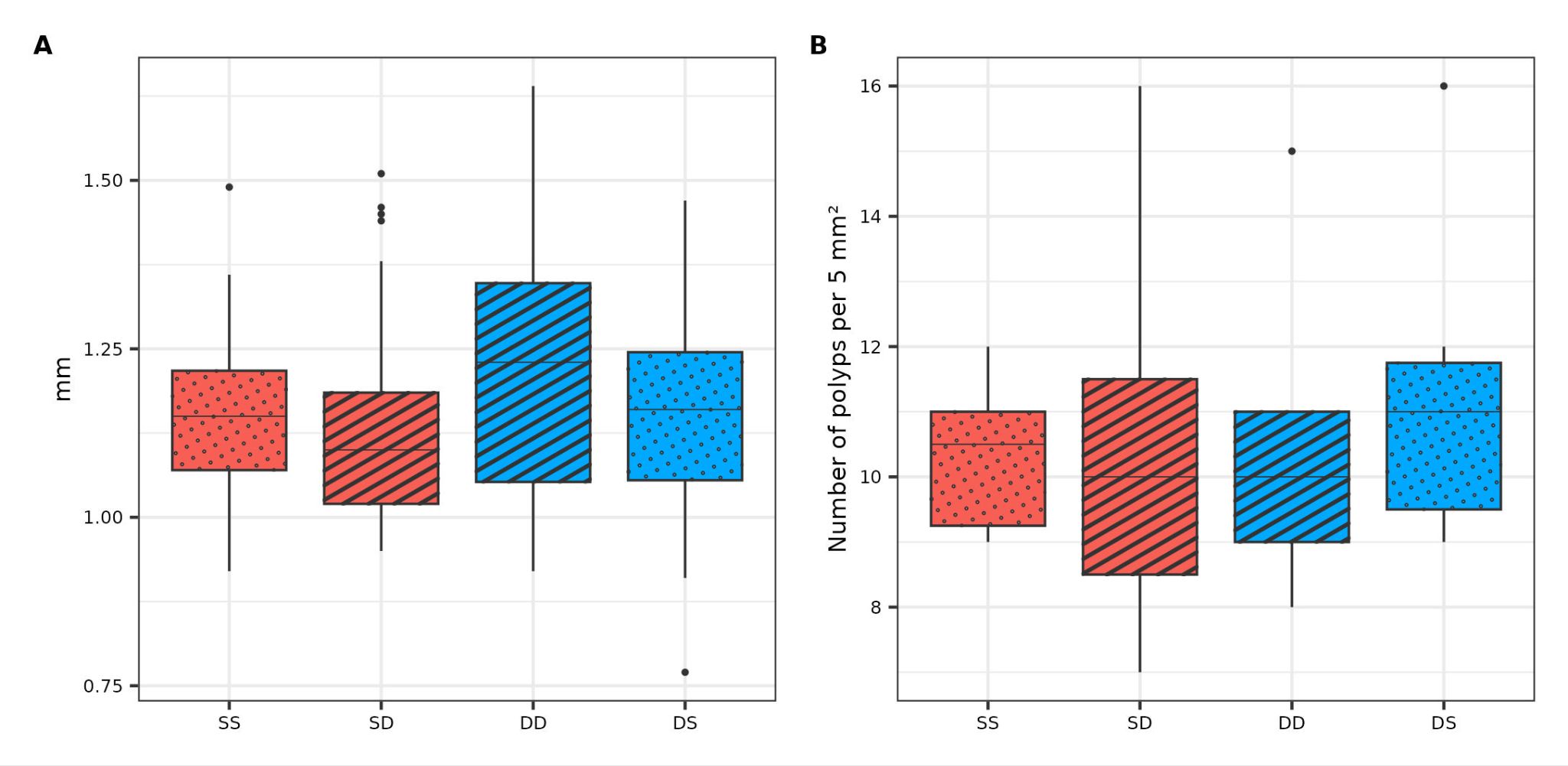
**Fig. S5.** Corallite size and density across transplantation treatments. Boxes show median and interquartile range; whiskers indicate 1.5× interquartile range; points represent outlier samples. **(A)** Corallite diameter. **(B)** Polyp count per 5 mm².

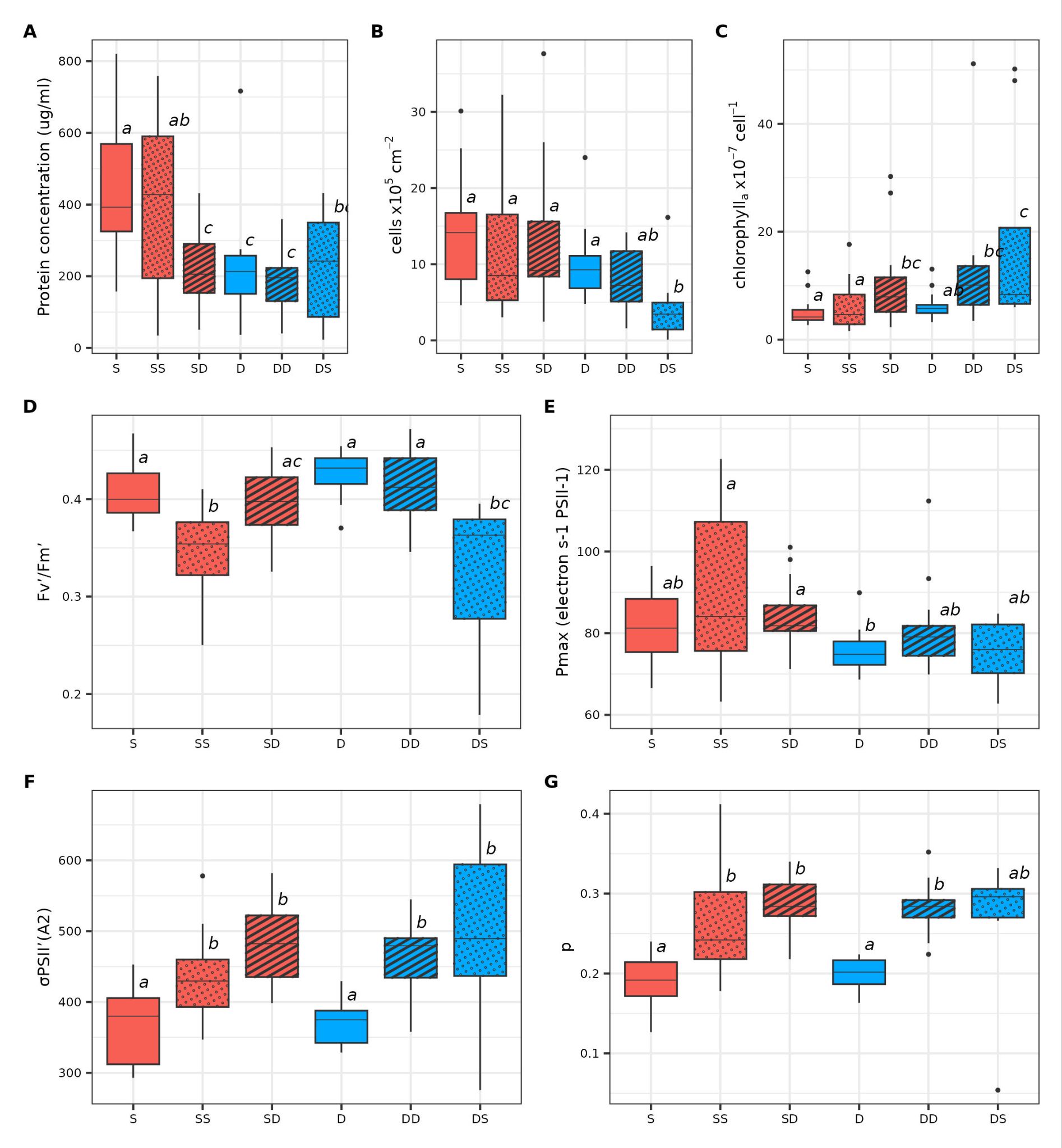

**Fig. S6.** Physiological responses of coral hosts and algal symbionts across experimental groups. Boxes show median and interquartile range; whiskers indicate 1.5× interquartile range; points represent outlier samples. Different lowercase letters denote statistically significant differences among treatments within each panel (p < 0.05). See Table S4 for test statistics. **(A)** Coral host protein concentration. **(B)** Symbiont cell density normalized to coral surface area. **(C)** Chlorophyll content per algal cell. **(D)** Maximum quantum yield of photosystem II (Fv/Fm). **(E)** Maximum photosynthetic rate (Pmax). **(F)** Functional absorption cross-section of PSII (σPSII). **(G)** Connectivity parameter (p).

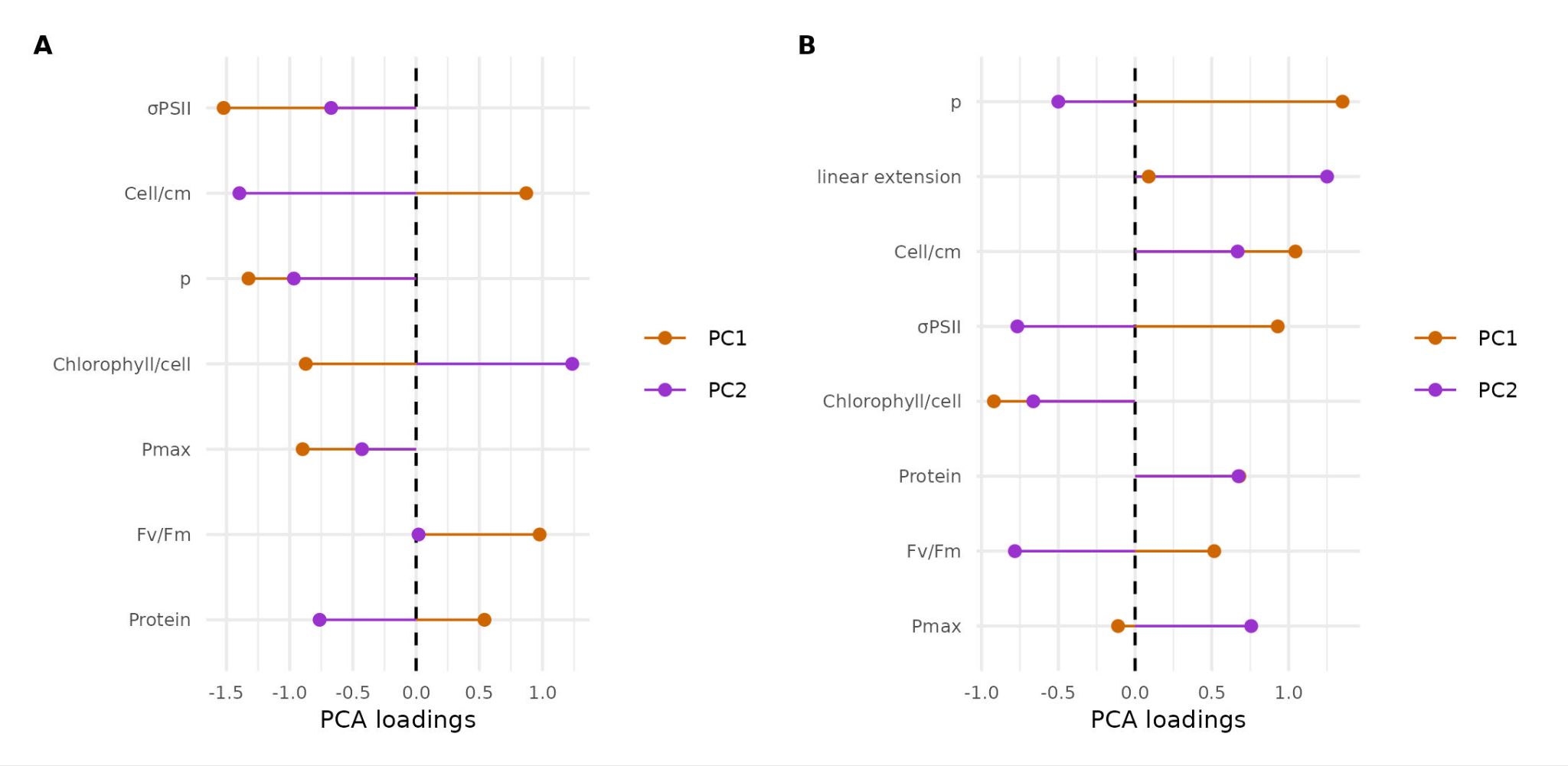
**Fig. S7.** Principal component loadings of coral traits. **(A)** PC1 and PC2 loadings for the variation of the coral physiological traits measured pre- and post-translocation. **(B)** PC1 and PC2 loadings for the variation of the coral physiological and skeletal traits measured post-translocation. Loadings indicate the strength and direction of the correlation between each trait and principal component.

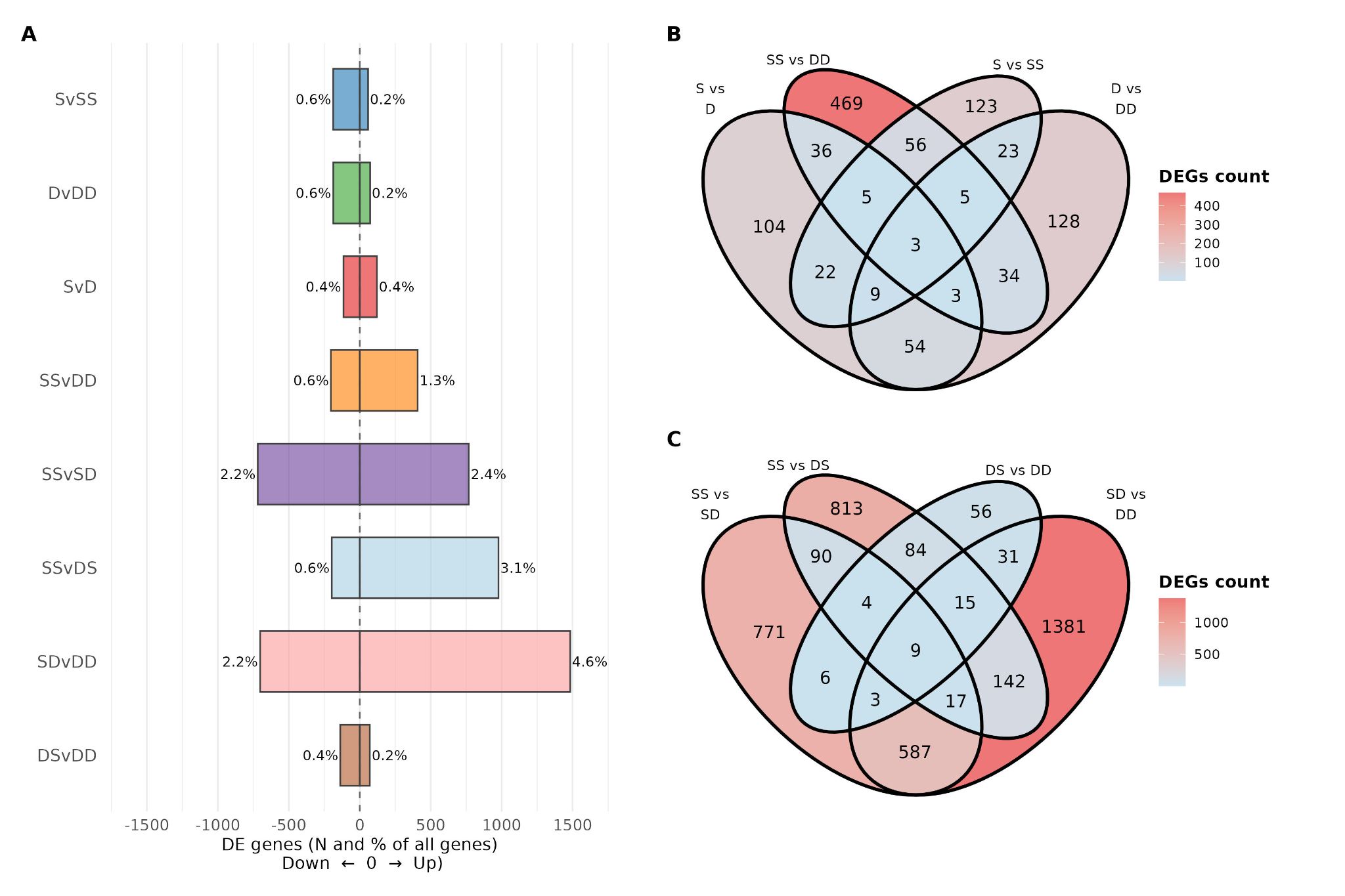

**Fig. S8.** DEG abundance and overlap across groups. (A) Percentage and number of DEGs (adjusted p < 0.05) for each contrast. Bars indicate up- and downregulated genes. (B) Venn diagram illustrating the overlap of DEGs among native groups. (C) Venn diagram illustrating the overlap of DEGs among translocation groups.

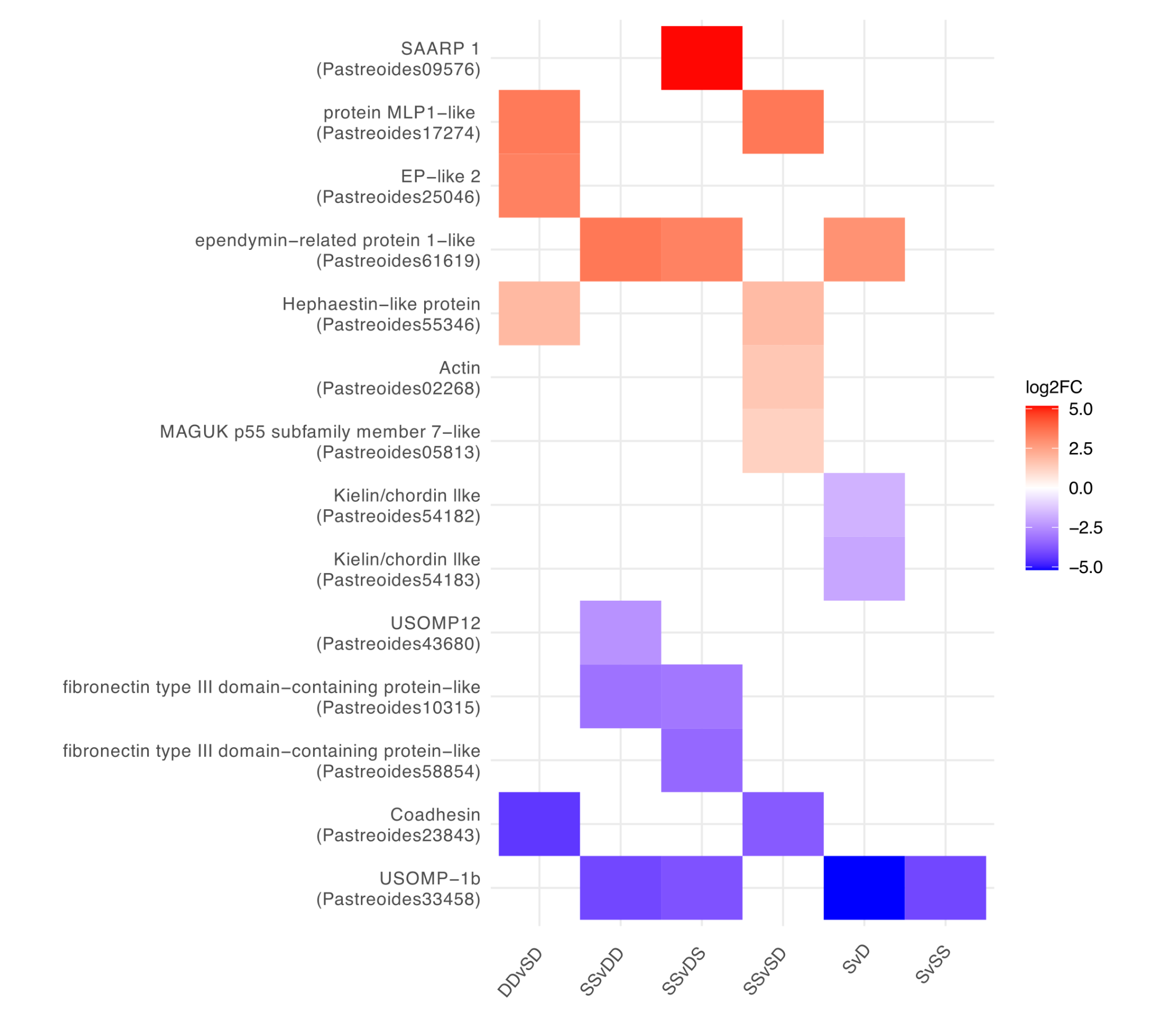
**Fig. S9.** Heatmap of differentially expressed biomineralization-related genes. Tile color represents log2 fold change values for coral biomineralization-related genes identified by orthology. Positive values indicate higher expression in the first group of each contrast. Contrasts without differentially expressed biomineralization genes are not shown.

| **site** | **depth** | **max_PAR** | **min_PAR** | **mean_PAR** | **sd_PAR** | **se_PAR** | **mean_delta_PAR** | **sd_delta_PAR** | **se_delta_PAR** |
| --- | --- | --- | --- | --- | --- | --- | --- | --- | --- |
| **Coral City** | 10 | 756.2 | 13.3 | 435.4 | 169 | 15.7 | 502.5 | 101.6 | 45.5 |
| **Coral City** | 40 | 190.9 | 2 | 92.8 | 42.8 | 4 | 113.9 | 40.2 | 18 |
| **Martha’s Finyard** | 10 | 745.4 | 2.9 | 446.6 | 170.1 | 12.3 | 556.4 | 67.1 | 23.7 |
| **Martha’s Finyard** | 40 | 302.7 | 1.2 | 171.2 | 71.5 | 5.1 | 244.6 | 17.3 | 6.1 |

**Table S1.** Descriptive statistics of midday (12:00 – 14:00) average photosynthetically active radiation (PAR) recorded by in situ loggers at two reef sites (Coral City and Martha’s Finyard) and two depths (10 m and 40 m). For each site–depth combination, the table reports maximum, minimum, mean, standard deviation (sd), and standard error (se) of PAR, as well as the mean monthly PAR range (mean ΔPAR) with associated sd and se. PAR is expressed in μmol photons m⁻² s⁻¹.

| **site** | **depth** | **min_temp** | **max_temp** | **mean_temp_val** | **sd_temp** | **se_temp** | **mean_delta_temp** | **sd_delta_temp** | **se_delta_temp** |
| --- | --- | --- | --- | --- | --- | --- | --- | --- | --- |
| **Coral City** | 10 | 27 | 30.98 | 29.03 | 1.03 | 0.1 | 1.02 | 0.42 | 0.19 |
| **Coral City** | 40 | 27.05 | 30.42 | 28.12 | 0.85 | 0.08 | 1.03 | 0.47 | 0.21 |
| **Martha’s Finyard** | 10 | 27.08 | 30.55 | 29.26 | 0.94 | 0.07 | 0.92 | 0.39 | 0.14 |
| **Martha’s Finyard** | 40 | 27.06 | 30.17 | 28.8 | 0.97 | 0.07 | 0.81 | 0.49 | 0.17 |

**Table S2.** Descriptive statistics of daily average temperature recorded by in situ loggers at two reef sites (Coral City and Martha’s Finyard) and two depths (10 m and 40 m). For each site–depth combination, the table reports maximum, minimum, mean, standard deviation (sd), and standard error (se) of temperature, as well as the mean monthly temperature range (mean Δtemperature) with associated sd and se. Temperature is expressed in °C.

| **Feature** | **Statistical test** | **Test value** | ***p value*** | **n** | **Post hoc test** |
| --- | --- | --- | --- | --- | --- |
| Coral host protein concentration | LMM (Linear mixed model) | F(5, 86.54) = 8.03 | 2.962e-06 | 97 | Tukey-adjusted |
| Symbiont cell density per surface area | LMM | F₍5,91₎ = 5.03 | 0.0004121 | 97 | Tukey-adjusted |
| Chlorophyll content per algal cell | GLMM (Generalized linear mixed model, inverse Gaussian, log link) | *χ*²(5) = 32.523 | 4.68e-06 | 97 | Tukey-adjusted |
| Functional absorption cross-section of PSII (σPSII). | LMM with fitted group-specific residual variance | F₍5,57₎ = 21.62 | <0.0001 | 99 | Tukey-adjusted |
| Maximum quantum yield of photosystem II (Fv/Fm) | LMM with fitted group-specific residual variance | F₍5,57₎ = 16.26 | <0.0001 | 99 | Tukey-adjusted |
| Maximum photosynthetic rate (Pmax) | LMM with fitted group-specific residual variance | F₍5,57₎ = 4.41 | 0.0018 | 99 | Tukey-adjusted |
| Connectivity parameter (p) | LMM with fitted group-specific residual variance | F₍5,57₎ = 67.06 | <0.0001 | 99 | Tukey-adjusted |
| Skeletal linear extension | LMM | F₍3,39.72₎ = 15.13 | 1.022e-06 | 60 | Tukey-adjusted |
| Alizarin band thickness | LMM | F₍3,57₎ = 10.74 | 1.124e-05 | 60 | Tukey-adjusted |
| Corallite diameter | LMM | F₍3,20.1₎ = 0.76 | 0.5312 | 161 |  |
| Corallite density | Poisson GLM (log link) | *χ*² = 0.33 | 0.9524 | 24 |  |

**Table S3.** Summary of statistical tests for physiological and skeletal traits across treatment groups.

| **PCA analysis** | **Parameter** | **PC1** | **PC2** | **PC3** | **R²** | **p-value** |
| --- | --- | --- | --- | --- | --- | --- |
| **Pre- and post-translocation** | Temperature | -0.018 | 0.087 | -0.339 | 0.12 | 0.005 |
|  | PAR | 0.242 | 0.029 | -0.411 | 0.23 | 0.001 |
| **Post-translocation only** | Temperature | −0.211 | 0.509 | -0.081 | 0.31 | 0.002 |
|  | PAR | −0.118 | 0.718 | -0.206 | 0.57 | 0.001 |

**Table S4.** Loadings, coefficients of determination (R²), and permutation-based p-values for environmental vectors fitted to PCA. The first PCA includes physiological variables with environmental data measured pre- and post-translocation. The second PCA includes physiological and skeletal variables using environmental measurements collected post-translocation only. PAR = photosynthetically active radiation.

| **Dataset** | **Source** | **Df** | **Sum of squares** | **R²** | **F** | **p-value** |
| --- | --- | --- | --- | --- | --- | --- |
| **Pre- and post-translocation** | Treatment | 5 | 201.96 | 0.314 | 7.95 | 0.001 |
|  | Residual | 87 | 442.04 | 0.686 | — | — |
|  | Total | 92 | 644.00 | 1.000 | — | — |
| **Post-translocation only** | Origin depth | 1 | 31.89 | 0.071 | 4.75 | 0.001 |
|  | Transplant depth | 1 | 42.74 | 0.095 | 6.365 | 0.001 |
|  | Origin depth:Transplant depth | 1 | 17.55 | 0.039 | 2.614 | 0.021 |
|  | Residual | 53 | 355.82 | 0.794 | — | — |
|  | Total | 56 | 448 | 1.000 | — | — |

**Table S6.** PERMANOVA results (Euclidean distance, 999 permutations) testing for treatment effects on multivariate trait space. Analyses were performed separately for (i) physiological traits using measurements collected pre- and post-translocation and (ii) physiological and skeletal traits using post-translocation data only. R² indicates the proportion of variance explained by treatment.

| **Dataset** | **Comparison** | **Sum of squares** | **F** | **R²** | **p-value** | **p-adjusted** |
| --- | --- | --- | --- | --- | --- | --- |
| **Pre- and post-translocation** | **S vs DD** | 69.140 | 22.502 | 0.391 | 0.001 | 0.015 |
|  | **S vs D** | 18.308 | 6.890 | 0.169 | 0.001 | 0.015 |
|  | **S vs SD** | 70.451 | 19.522 | 0.352 | 0.001 | 0.015 |
|  | **S vs DS** | 60.180 | 8.135 | 0.253 | 0.001 | 0.015 |
|  | **S vs SS** | 35.904 | 6.958 | 0.170 | 0.001 | 0.015 |
|  | **DD vs D** | 35.651 | 14.077 | 0.312 | 0.001 | 0.015 |
|  | **DD vs SS** | 43.636 | 8.268 | 0.211 | 0.001 | 0.015 |
|  | **D vs SD** | 49.454 | 15.687 | 0.329 | 0.001 | 0.015 |
|  | **D vs DS** | 49.405 | 6.653 | 0.250 | 0.001 | 0.015 |
|  | **D vs SS** | 51.961 | 10.647 | 0.262 | 0.001 | 0.015 |
|  | **SD vs SS** | 26.187 | 4.506 | 0.123 | 0.001 | 0.015 |
| **Post-translocation only** | **DD vs SS** | 58.663 | 10.097 | 0.246 | 0.001 | 0.006 |
|  | **SD vs SS** | 39.397 | 6.53 | 0.169 | 0.001 | 0.006 |

**Table S7.** Pairwise PERMANOVA results for (i) physiological traits using measurements collected pre- and post-translocation and (ii) physiological and skeletal traits using post-translocation data only based on Euclidean distances among multivariate trait measurements. Analyses were conducted using the pairwise.adonis function with 999 permutations. Reported values include pseudo-F statistics, R², and permutation-based p-values adjusted for multiple testing using Bonferroni correction. Only significant comparisons (p-adjusted < 0.05) are shown.

| **GOSLIM** | **GOSLIM_name** | **direction** | **n_studies** | **n_total** | **per_study_counts** |
| --- | --- | --- | --- | --- | --- |
| GO:0048856 | anatomical structure development | down | 3 | 13 | CurrentStudy:1; M2020:7; Sc2023b:5 |
| GO:0016787 | hydrolase activity | down | 2 | 11 | CurrentStudy:1; Sc2023b:10 |
| GO:0022414 | reproductive process | down | 2 | 9 | M2020:1; Sc2023b:8 |
| GO:0006629 | lipid metabolic process | down | 3 | 8 | Sc2023a:1; Sc2023b:6; SV2020:1 |
| GO:0030154 | cell differentiation | down | 2 | 6 | M2020:2; Sc2023b:4 |
| GO:0007155 | cell adhesion | down | 3 | 5 | M2020:3; Sc2023a:1; Sc2023b:1 |
| GO:0005215 | transporter activity | down | 2 | 4 | CurrentStudy:1; Sc2023b:3 |
| GO:0016740 | transferase activity | down | 2 | 4 | M2020:1; Sc2023b:3 |
| GO:0098772 | molecular function regulator activity | down | 2 | 4 | CurrentStudy:1; Sc2023a:3 |
| GO:0007018 | microtubule-based movement | down | 2 | 3 | M2020:2; Sc2023b:1 |
| GO:0016853 | isomerase activity | down | 2 | 3 | M2020:1; Sc2023b:2 |
| GO:0006355 | regulation of DNA-templated transcription | down | 2 | 2 | Sc2023a:1; Sc2023b:1 |
| GO:0016740 | transferase activity | up | 3 | 30 | CurrentStudy:1; Sc2023b:28; T2021:1 |
| GO:0065003 | protein-containing complex assembly | up | 2 | 30 | CurrentStudy:1; Sc2023b:29 |
| GO:0098772 | molecular function regulator activity | up | 3 | 28 | Sc2023a:1; Sc2023b:25; T2021:2 |
| GO:0055086 | nucleobase-containing small molecule metabolic process | up | 3 | 21 | M2020:1; Sc2023b:19; T2021:1 |
| GO:0002376 | immune system process | up | 2 | 21 | Sc2023a:1; Sc2023b:20 |
| GO:0016787 | hydrolase activity | up | 2 | 21 | M2020:3; Sc2023b:18 |
| GO:0016491 | oxidoreductase activity | up | 3 | 15 | CurrentStudy:1; M2020:1; Sc2023b:13 |
| GO:0016829 | lyase activity | up | 3 | 10 | M2020:1; Sc2023b:6; T2021:3 |
| GO:0005215 | transporter activity | up | 2 | 8 | CurrentStudy:1; Sc2023b:7 |
| GO:0016071 | mRNA metabolic process | up | 2 | 8 | Sc2023b:7; SV2020:1 |
| GO:0006091 | generation of precursor metabolites and energy | up | 2 | 7 | CurrentStudy:3; Sc2023b:4 |
| GO:0006886 | intracellular protein transport | up | 2 | 6 | Sc2023b:5; SV2020:1 |
| GO:0006575 | modified amino acid metabolic process | up | 2 | 4 | CurrentStudy:1; Sc2023b:3 |
| GO:0140110 | transcription regulator activity | up | 2 | 3 | M2020:1; Sc2023b:2 |

**Table S7.** Summary of GO slim functional categories used to construct the combined word cloud of recurrent transcriptional responses (Fig. 7). The table lists GO-slim IDs (GOSLIM) and term names (GOSLIM_name) detected across studies, separated by enrichment direction (up or down). For each term, the number of independent studies in which it occurred (n_studies; reflects word intensity), the total number of occurrences across all studies (n_total; used to scale word size), and the per-study occurrence counts are provided. Only GO-slim terms detected in at least two independent studies are included. The underlying dataset of GO terms and GO slims across these studies can be found in Data S4.

| **Sample** | **Raw reads, million pairs** | **Trimmed reads, million pairs** | **rRNA %** | **Host-mapped reads, million** | **Gene-level counts** |
| --- | --- | --- | --- | --- | --- |
| **100-CC-SD-73** | 34.7 | 27.3 | 42.2 | 4.8 | 1,127,535 |
| **102-CC-DD-82** | 33.5 | 29.1 | 38.1 | 3.9 | 2,078,265 |
| **108-CC-DD-94** | 34.3 | 28.7 | 67.9 | 4.2 | 1,779,893 |
| **114-MF-SD-45** | 41.4 | 33.7 | 14.0 | 16.7 | 6,716,295 |
| **117-MF-SD-44** | 29.6 | 25 | 4.6 | 15.1 | 6,026,051 |
| **123-CC-DD-81** | 34.6 | 30.1 | 3.1 | 21.1 | 11,922,704 |
| **23-MF10-1a** | 35.3 | 31.1 | 31.9 | 15.4 | 8,781,589 |
| **25-CC40-2b** | 31.8 | 27.5 | 3.1 | 20.7 | 12,056,044 |
| **27-MF40-2b** | 32.1 | 27.8 | 54.5 | 5.4 | 2,954,261 |
| **28-CC10-9b** | 31.7 | 27.5 | 57.0 | 8.4 | 4,990,476 |
| **31-MF40-5b** | 44.2 | 38.4 | 83.6 | 5.6 | 2,620,263 |
| **32-CC10-1b** | 30.5 | 26.5 | 86.6 | 3 | 1,259,411 |
| **35-MF40-1b** | 31.3 | 27.3 | 45.4 | 11.6 | 6,884,284 |
| **37-CC40-8b** | 38.3 | 34 | 71.0 | 8.4 | 4,687,437 |
| **4-MF10-4a** | 31.7 | 26.4 | 58.4 | 8.5 | 4,931,598 |
| **42-MF10-10b** | 37.9 | 33.4 | 21.1 | 20.3 | 12,114,648 |
| **48-CC10-6b** | 27.8 | 24 | 86.7 | 2.8 | 866,868 |
| **49-CC40-1b** | 26.3 | 23.4 | 62.3 | 5.8 | 3,313,338 |
| **63-MF-SS-60** | 38.1 | 32.6 | 19.2 | 20.1 | 11,733,448 |
| **66-MF-DS-52** | 28.5 | 20.6 | 79.4 | 2.8 | 1,199,588 |
| **73-CC-SS-38** | 45 | 36.2 | 79.4 | 5.5 | 2,622,399 |
| **75-CC-DD-84** | 31.1 | 26.6 | 20.9 | 12.2 | 7,256,215 |
| **78-MF-DS-54** | 36.7 | 14.7 | 62.4 | 2.6 | 599,639 |
| **80-1-CC-DS-86** | 30.2 | 25.9 | 2.2 | 19.4 | 11,845,253 |
| **81-CC-DS-87** | 35 | 21.7 | 51.8 | 7.6 | 4,194,143 |
| **84-CC-DD-80** | 36.7 | 20.8 | 71.8 | 4.2 | 2,146,775 |
| **91-CC-DS-89** | 20.3 | 69.5 | 87.1 | 1.4 | 535,645 |
| **94-CC-SS-37** | 30.9 | 26.2 | 73.5 | 4.6 | 2,322,927 |
| **97-MF-SS-64** | 29.3 | 22.5 | 40.3 | 3 | 852,965 |
| **98-MF-SS-69** | 35.1 | 29.2 | 64.1 | 7.3 | 3,822,419 |
| **99-MF-SD-43** | 30.9 | 27.1 | 2.0 | 13.4 | 4,909,598 |
| **mean** | 33.4 | 28.9 |  | 9.2 | 4,811,354 |
| **median** | 32.1 | 27.3 |  | 7.3 | 3,822,419 |

**Table S8.** Basic read statistics.

| **Link to the database** | **Symbiont species** | **Former Symbiodinium clade** |
| --- | --- | --- |
| <https://www.ncbi.nlm.nih.gov/datasets/genome/GCA_963970005.1/> | *Durusdinium trenchii* | former clade D |
| <https://marinegenomics.oist.jp/symb/viewer/info?project_id=21> | *Breviolum minutum* | former clade B |
| <https://marinegenomics.oist.jp/symb/viewer/download?project_id=37> | *Symbiodinium clade A3* |  |
| <https://espace.library.uq.edu.au/view/UQ:f1b3a11> | *Symbiodinium tridacnidorum* | former clade A |
| <https://www.ncbi.nlm.nih.gov/datasets/genome/GCA_905231905.1/> | *Symbiodinium pilosum* | former clade A |
| <https://espace.library.uq.edu.au/view/UQ:f1b3a11> | *Symbiodinium necroappetens* | former clade A |
| <https://espace.library.uq.edu.au/view/UQ:f1b3a11> | *Symbiodinium natans* | former clade A |
| <http://smic.reefgenomics.org/> | *Symbiodinium microadriaticum* | former clade A |
| <https://espace.library.uq.edu.au/view/UQ:f1b3a11> | *Symbiodinium linucheae* | former clade A |
| <http://symbs.reefgenomics.org/download/> | *Fugacium kawagutii* | former clade F |
| <http://symbs.reefgenomics.org/download/> | *Cladocopium goreau* | former clade C1 |
| <https://marinegenomics.oist.jp/symb/viewer/download?project_id=40> | *Cladocopium C15* |  |

**Table S9.** Symbiodiniaceae proteomes used in symbiont composition analysis

**Data S1. (separate file)**

Gene Ontology enrichment results for differentially expressed genes across *Porites astreoides* experimental groups. GO term enrichment was performed using the full set of differentially expressed genes for each contrast, without separating up- and downregulated genes. Columns are defined as follows: ID, GO term identifier; Description, GO term name; GeneRatio, proportion of input genes associated with the term; BgRatio, proportion of background genes associated with the term; RichFactor, ratio of observed to background annotation frequency; FoldEnrichment, enrichment of the term relative to background expectation; zScore, direction and magnitude of enrichment based on up/down gene balance; pvalue, raw enrichment p value; p.adjust, multiple-testing–adjusted p value; qvalue, false discovery rate estimate; geneID, list of overlapping genes; Count, number of overlapping genes contributing to the term. Only GO terms with p.adjusted<0.05 were retained. The contrasts with no significant GO terms are not shown.

**Data S2. (separate file)**

Gene Ontology enrichment results for direction-specific gene sets across experimental groups. GO term enrichment was performed separately for upregulated and downregulated gene sets for each contrast. Columns are defined as follows: ID, GO term identifier; Description, GO term name; GeneRatio, proportion of input genes associated with the term; BgRatio, proportion of background genes associated with the term; RichFactor, ratio of observed to background annotation frequency; FoldEnrichment, enrichment of the term relative to background expectation; zScore, direction and magnitude of enrichment based on up/down gene balance; pvalue, raw enrichment p value; p.adjust, multiple-testing–adjusted p value; qvalue, false discovery rate estimate; geneID, list of overlapping genes; Count, number of overlapping genes contributing to the term. Only GO terms with p.adjusted<0.05 were retained. The contrasts with no significant GO terms are not shown.

**Data S3. (separate file)**

Gene Ontology enrichment results for coexpression module clusters. Gene Ontology (GO) term enrichment analysis of genes showing significant module–trait correlations within each coexpression module cluster. Columns are defined as follows: ID, GO term identifier; Description, GO term name; GeneRatio, proportion of input genes associated with the term; BgRatio, proportion of background genes associated with the term; RichFactor, ratio of observed to background annotation frequency; FoldEnrichment, enrichment of the term relative to background expectation; zScore, direction and magnitude of enrichment based on up/down gene balance; pvalue, raw enrichment p value; p.adjust, multiple-testing–adjusted p value; qvalue, false discovery rate estimate; geneID, list of overlapping genes; Count, number of overlapping genes contributing to the term. Only GO terms with p.adjusted<0.05 were retained. The clusters with no significant GO terms are not shown.

**Data S4. (separate file)**

Differentially expressed Gene Ontology (GO) terms compiled across transcriptomic studies included in the comparative analysis. Each row corresponds to a reported GO term and includes the following information: study identifier (study_id), GO identifier (GO_ID) and term name (GO_term), direction of enrichment (up or down), reported significance value (p.value), source environment (description), study species (species), source publication DOI (DOI), and the corresponding GO-slim category and term name (GOSLIM, GOSLIM_name). This dataset served as a basis for generating Table S7 and Fig. 7.
